# Synergy in viral-bacterial coinfection expedites algal bloom demise

**DOI:** 10.1101/2025.08.04.668390

**Authors:** Eliran Soffer, Constanze Kuhlisch, Jean-Baptiste Raina, Noa Barak-Gavish, Guy Tadmor, Guy Schleyer, Tal Sharon Nir-Zadock, Inbal Nussbaum, Estelle E. Clerc, Roman Stocker, Daniella Schatz, David Zeevi, Assaf Vardi

## Abstract

Algal blooms are hotspots for diverse microbial interactions affecting bloom dynamics. Viral infections often regulate the demise of these blooms, recycling >25% of oceanic fixed carbon. Although these blooms collapse synchronously, only 10-30% of algal cells show active viral infection, suggesting additional mortality agents. Using microbial communities from natural coccolithophore blooms, we discovered that bacteria significantly enhanced virus-mediated algal mortality. We quantified this interaction using two novel metrics: the Synergy Index (SI) measuring viral-bacterial interaction strength, and the Benefit Index (BI) assessing changes in bacterial fitness. We identified *Alteromonas* bacteria as drivers of viral-bacterial coinfection with consistently high SI and BI. In natural communities, *Alteromonas* exhibited chemoattraction towards infected algal cells, leading to their enrichment in bloom demise particles. This viral-bacterial synergy represents a previously unrecognized mortality agent controlling bloom termination, modulating the balance between the carbon-recycling viral shunt and the carbon-exporting viral shuttle pathways, likely influencing global carbon cycling.

## Introduction

About 50 gigatons of carbon are fixed annually by marine phototrophs, providing organic matter that feeds the entire microbial food web (*1–3*). A substantial proportion of this carbon fixation occurs during large-scale algal blooms (*4*, *5*). Therefore, microbial processes that control cell fate in these blooms can have profound impacts on carbon cycling in the ocean (*6*). Algal blooms are hotspots for diverse interactions and metabolic exchange with grazers, bacteria, fungi and viruses (*7*, *8*). As the most abundant biological entities in the ocean, viruses are major mortality agents driving nutrient cycling globally and significantly impacting algal bloom demise (*9*, *10*).

Viruses regulate the metabolic composition and fate of carbon and other nutrients (*11*) through two distinct ecosystem processes: the viral shunt, which fuels the microbial food web by catalyzing nutrient recycling in the surface ocean (*12*, *13*); and the viral shuttle, which facilitates carbon export through enhanced aggregation of lysed cells and sinking of particulate organic matter to the deep ocean (*13*, *14*). Viral lysis of algal populations creates a distinct chemical footprint (*15-17*), which can promote the growth of adapted bacteria, thereby altering bacterial biodiversity (*18*). Yet, how viral-induced shifts in microbial community composition influence algal cell fate – and thus global carbon cycling – remains poorly understood (*19*).

We selected *Gephyrocapsa huxleyi* (formerly *Emiliania huxleyi*) to study algae-virus-bacteria interactions due to its ecological significance in marine ecosystems. This cosmopolitan unicellular eukaryotic alga forms extensive oceanic blooms, covering thousands of square kilometers (*20*). It plays a crucial role in global carbon and sulfur biogeochemical cycles due to its photosynthetic ability, calcite exoskeleton production and production of sulfur volatiles such as dimethyl sulfide (DMS) (*20–22*). These blooms are often terminated by a giant dsDNA virus, the *E. huxleyi* virus (EhV; (*23*)). Recent quantification of active viral infection in the marine environment using novel single-cell approaches (*24*, *25*) revealed a puzzling phenomenon: less than 30% of algal cells were actively infected by the virus, despite the synchronized collapse of entire blooms (*26*). This discrepancy suggests that additional mortality agents may act in concert with viral infection during the demise of algal blooms. Inspired by findings from the biomedical field demonstrating that viral-bacterial coinfections can synergistically increase mortality (*27*, *28*), we investigated the interplay between bacteria and viruses during virus-induced algal bloom demise. Using a bacterial consortium isolated from natural algal blooms (**Fig. S1**), we show that bacteria can exacerbate virus-induced algal mortality, revealing a previously unrecognized tripartite interaction with potentially significant implications for global biogeochemical cycles.

## Results

### Natural algae-associated bacteria expedite algal death upon viral infection

We sought to determine the role of bacteria as additional mortality agents during virus-induced algal bloom demise. We hypothesized that if bacteria naturally associated with an alga contribute to its mortality, then an antibiotic treatment would reduce the overall death rate during viral infection. To test this hypothesis, we cultured a xenic strain of *G. huxleyi* isolated as a single cell from an induced bloom during a mesocosm experiment in a fjord near Bergen, Norway (*18*). In the absence of viral infection, the xenic culture showed comparable algal growth patterns regardless of antibiotic treatment (Two-way ANOVA, p > 0.05; **Fig. 1a**, blue lines), suggesting that the bacteria naturally associated with *G. huxleyi* maintain a commensal or mutualistic relationship rather than a pathogenic one under normal growth conditions.

**Figure 1.**
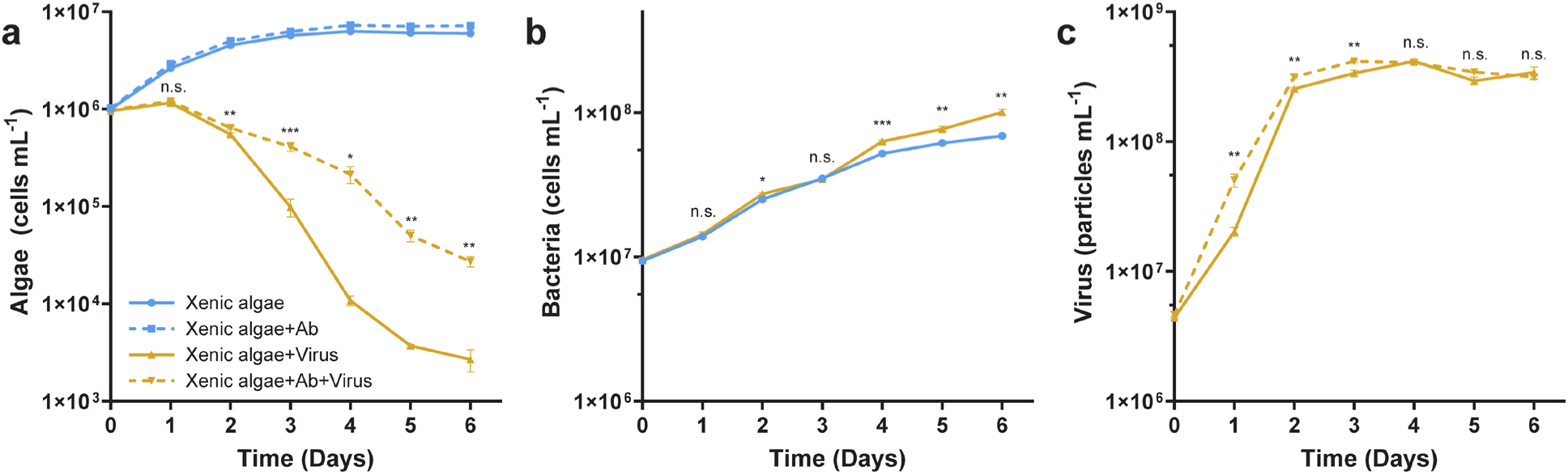
*G. huxleyi*-associated bacteria synergistically enhance virus-induced algal mortality. (a) Growth curves of xenic *G. huxleyi* under four conditions: uninfected with/without antibiotics (blue lines) and EhV-infected with/without antibiotics (yellow lines). (b) Bacterial abundance during cultivation with uninfected or EhV-infected *G. huxleyi*. (c) Viral production in antibiotics-treated and untreated cultures. * p < 0.05, ** p < 0.01, *** p < 0.001, **** p < 10^-4^; statistical significance determined by two-way ANOVA with Sídák’s post-hoc test; n = 4. Ab, antibiotics.

Infection with EhV induced algal population decline from day 1 (**Fig. 1a**). However, when antibiotics were administered to reduce the abundance of the algal microbiome (**Fig. S2**), we observed reduced algal mortality compared to the untreated xenic culture (Two-way ANOVA, p < 0.01 from day 2; **Fig. 1a**, yellow lines). This difference demonstrates that the bacterial community may contribute significantly to algal mortality, but only during viral infection, indicating a potential synergistic pathogenic interaction between viruses and bacteria. Analysis of bacterial abundances revealed that in the xenic culture during viral infection, bacteria grew to higher abundances compared to the uninfected condition (Two-way ANOVA, p < 0.01 from day 4; **Fig. 1b**). Importantly, viral production remained unchanged between antibiotic-treated and untreated algal cultures when the difference in algal mortality was the highest (Two-way ANOVA, p > 0.05 from day 4; **Fig. 1c**).

Based on these findings, we propose that bacteria and viruses exhibit synergistic killing, whereby viral infection creates favorable conditions for bacterial pathogenicity, resulting in accelerated algal mortality that benefits bacterial growth. These results reveal that bacteria naturally associated with *G. huxleyi* can expedite its demise during viral infection.

### Tripartite algae-virus-bacteria interactions expedite algal death

We sought to disentangle the complex interactions between algae, viruses and bacteria during algal bloom demise. To this end, we developed a tripartite laboratory model system using algae, virus and bacteria strains that were isolated from algal blooms (**Fig. S1**). We isolated the bloom-associated microbiome (BAM) from the demise phase and determined its impact on *G. huxleyi* during viral infection.

When we reconstituted the tripartite system by exposing axenic *G. huxleyi* cultures to EhV with and without the BAM community, we observed an enhancement of virus-induced mortality in the presence of bacteria (**Fig. 2a**). While viral infection alone significantly reduced algal abundance compared to uninfected cultures, the addition of the BAM accelerated this decline, resulting in approximately 4-fold lower algal cell densities 7 days post infection (Two-way ANOVA, p < 10^-4^; **Fig. 2a**). This recapitulated our observations in xenic *G. huxleyi* cultures (**Fig. 1a**), confirming that bacterial presence synergistically enhances virus-mediated algal mortality.

**Figure 2.**
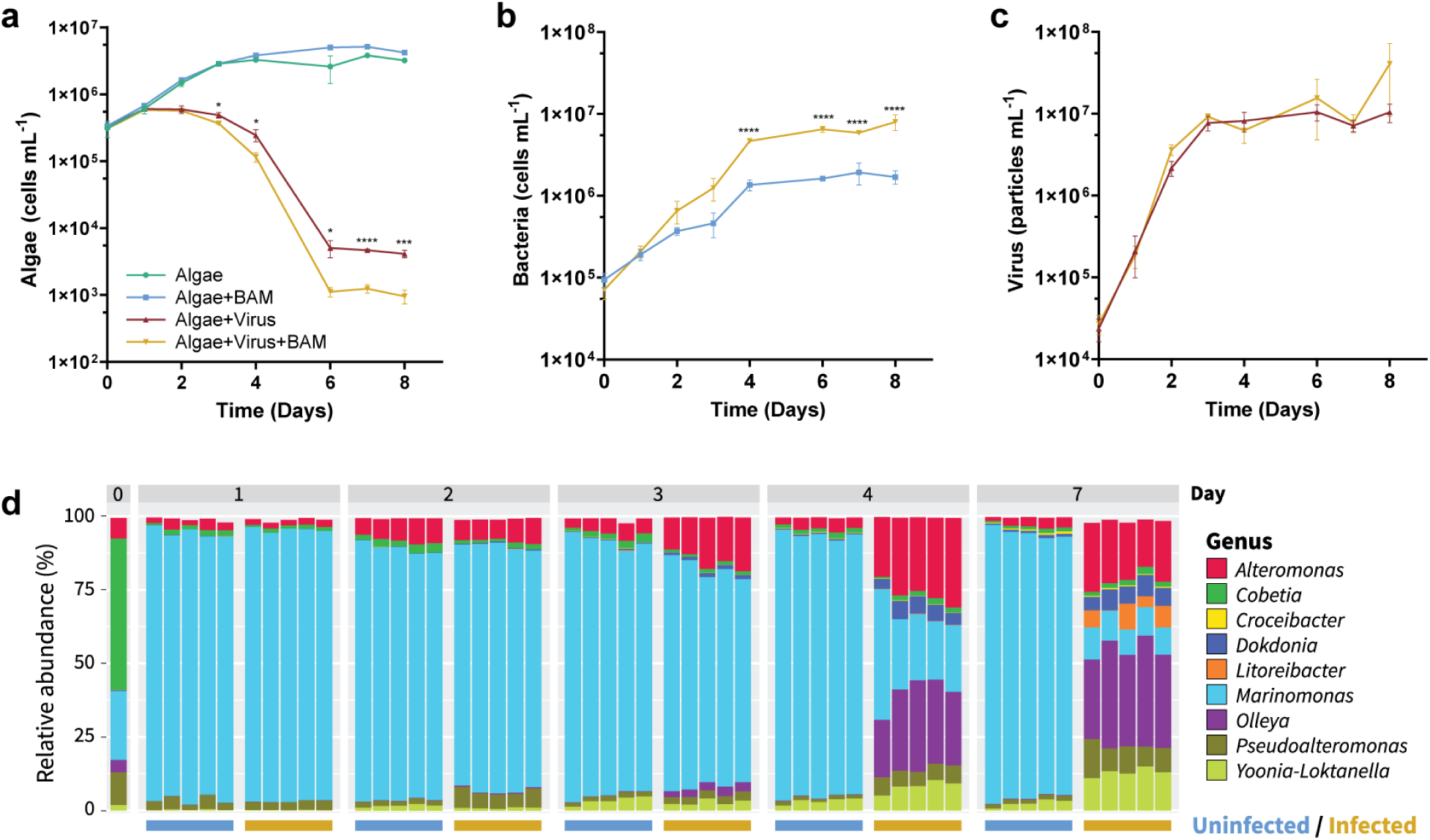
Tripartite interactions among algae, virus, and bacteria enhance algal mortality and reshape bacterial community composition. (a) Growth curves of axenic *G. huxleyi* under four conditions: uninfected, with bloom-associated microbiome (BAM) only, with EhV only, and with both BAM and EhV. (b) Bacterial abundances in cultures with and without viral infection. (c) Viral production in axenic and BAM-supplemented cultures. (d) Bacterial community composition over time in uninfected and EhV-infected cultures. * p < 0.05, ** p < 0.01, *** p < 0.001, **** p < 10^-4^; statistical significance determined by two-way ANOVA with Sídák’s post-hoc test; n = 5.

Bacterial population dynamics revealed significant proliferation in the tripartite system compared to the bacteria-only treatment (**Fig. 2b**). Similar to xenic *G. huxleyi* cultures (**Fig. 1b**), bacterial abundances in the virus-infected cultures were more than 4 times higher than in the uninfected cultures 8 days post infection (Two-way ANOVA, p < 10^-4^ from day 4), suggesting that bacterial communities exploit metabolites released during viral infection. Viral production remained comparable between axenic and BAM-supplemented cultures (Two-way ANOVA, p > 0.05; **Fig. 2c**), indicating that the bacterial consortium did not impair viral replication but rather complemented the viral killing mechanism.

To identify the microbial drivers of this tripartite interaction, we characterized changes in microbial community composition during viral infection using 16S rRNA amplicon sequencing (**Fig. 2d**). The bacterial community structure changed reproducibly in all replicates over the course of the viral infection. While the uninfected cultures maintained a relatively stable community dominated by *Marinomonas*, the virus-infected cultures showed a pronounced shift in community composition by day 4, coinciding with algal decline. Most notably, the genera *Alteromonas* and *Olleya* increased in relative abundance in infected cultures, rising from approximately 4.0±0.4% and 0.02±0.01%, respectively, at the beginning of the experiment to 20.2±2.7% and 32.9±4.4%, respectively, by day 7, thereby dominating the bacterial community (**Fig. 2d**). Concurrently, *Pseudoalteromonas* and *Yoonia-Loktanella* increased in their relative abundance, whereas *Marinomonas* declined. These findings suggest that viral infection creates a distinct metabolic niche that selectively promotes specific bacterial taxa capable of exploiting virus-induced changes in algal metabolism or cell lysis products.

### Quantitative metrics reveal distinct bacterial lifestyles during algal bloom demise

To systematically disentangle the microbial dynamics governing algal bloom demise, we developed two quantitative metrics to assess the contribution of different bacterial taxa to virus-induced algal mortality, and the effects of viral infection of algal cells on bacterial fitness. The Synergy Index (SI) quantifies the decrease in the algal population during viral infection in the presence of bacteria compared to viral infection alone. The Benefit Index (BI) measures the growth advantage that bacteria gain during viral infection compared to their growth with uninfected algae.

To calculate SI, we use the area under the growth curve of *G. huxleyi* during viral infection alone (AUC_virus_) and during bacterial and viral coinfection (AUC_tripartite_). We chose AUC as our metric because it provides a biologically meaningful measure of cumulative biomass change that captures complete temporal dynamics rather than single timepoints and offers robustness through multiple daily measurements. We calculated the difference between AUC_virus_ and AUC_tripartite_ and normalized this value using AUC_virus_, which is represented mathematically as 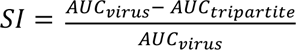 (**Fig. 3a**; **Methods**). Normalizing AUC differences creates a dimensionless metric enabling direct comparison across bacterial strains and experimental conditions regardless of baseline differences. A positive SI indicates that the bacterial presence reduces algal populations beyond that caused by viruses alone, with higher values reflecting stronger effects (e.g., SI = 10.0% means a 10% increase in tripartite-induced algal decline compared to mortality induced by viruses alone). When computing the SI for the effect of the bloom associated microbiome (BAM), we obtained a SI value equal to 18.2±0.8%, indicative of a substantial effect (**Fig. 3b**).

**Figure 3.**
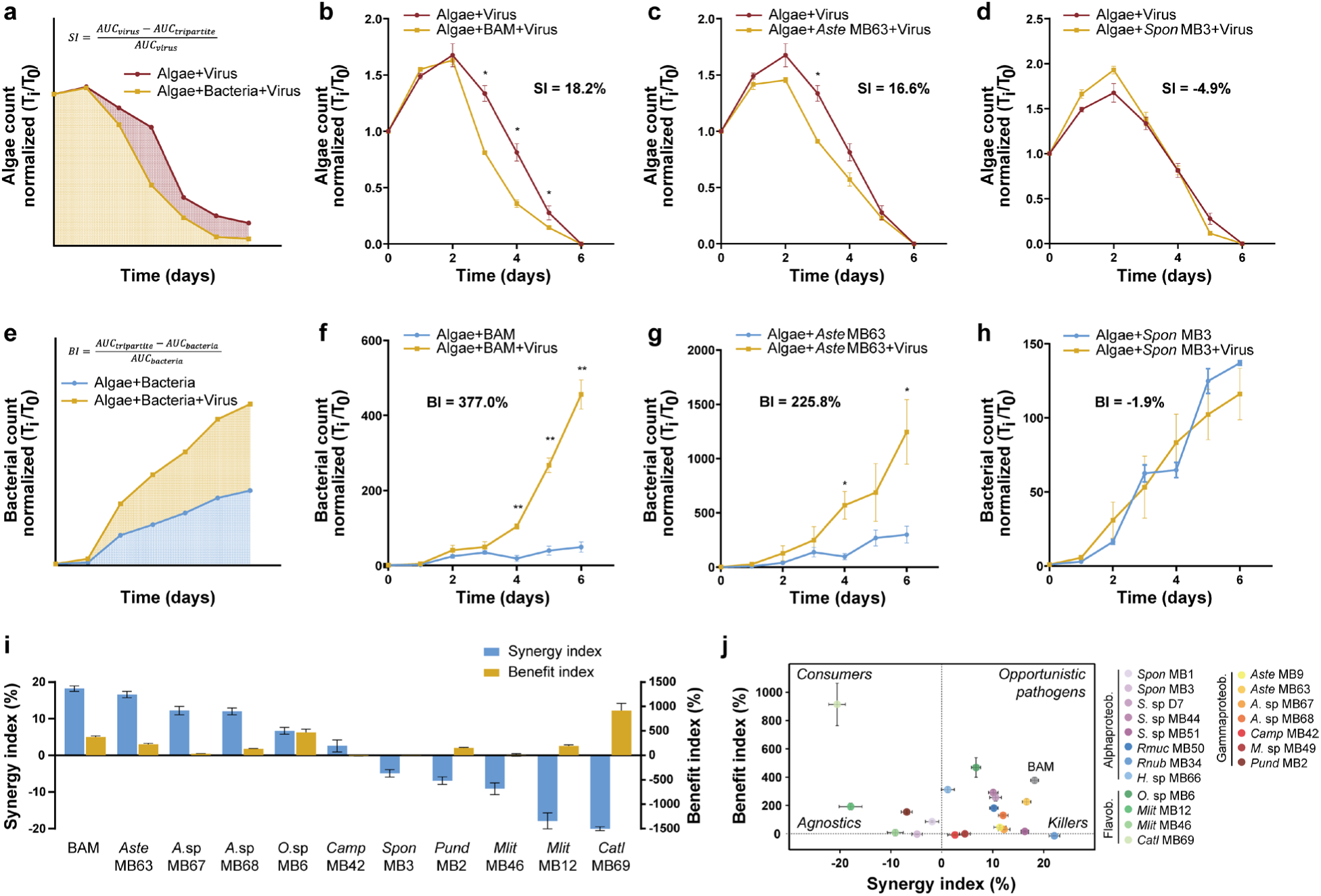
Quantitative metrics reveal species-specific bacterial ecological strategies in response to virus-induced algal mortality. (a) Schematic illustration of the Synergy Index (SI) calculation method comparing growth curves of virus-infected alga with or without bacteria. (b) Application of SI to assess the effect of the entire BAM community, showing significant enhancement of virus-induced mortality (SI = 18.2±0.8%). (c-d) Representative bacterial isolates from the BAM with contrasting SI values: *Alteromonas stellipolaris* MB63 (c) exhibits high synergistic killing (SI = 16.6±0.9%), while *Sulfitobacter pontiacus* MB3 (d) shows an absence of synergy (SI = -4.8±0.9%). (e) Schematic illustration of the Benefit Index (BI) calculation comparing bacterial growth curves in the presence of infected or uninfected alga. (f) Application of BI to the entire BAM community (BI = 337.0±22.4%), indicating substantial growth advantage during viral infection. (g-h) Bacterial growth curves showing contrasting BI values: *A. stellipolaris* MB63 (g) derives large benefit (BI = 225.8±23.4%) from coinfection, while *S. pontiacus* MB3 (h) shows no advantage (BI = -1.9±4.2%). (i) SI and BI values for ten isolated BAM members, revealing diverse ecological strategies. (j) Classification of bacterial ecological strategies for 19 algal bloom-associated bacteria based on SI and BI values, identifying opportunistic pathogens, killers, consumers, and agnostic bacteria. n = 3.

The Benefit Index (BI) quantifies the growth advantage bacteria derive from viral infection, calculated as 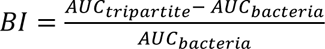, where *AUC_bacteria_* represents the area under the curve for bacterial growth with uninfected algae (**Fig. 3e**). The BAM community exhibited a high BI (377.0±22.4%), indicating that viral infection creates a metabolically favorable niche that enhances bacterial proliferation almost fourfold (**Fig. 3f**).

To identify key contributors to these synergistic interactions within the BAM, we examined the SI and BI for individual bacteria isolated from the BAM (**Table S1**). *Alteromonas stellipolaris* MB63 demonstrated a high SI (16.6±0.9%), indicating that this species enhanced virus-induced algal population decline by nearly 17% (**Fig. 3c**). In contrast, *Sulfitobacter pontiacus* MB3 showed a small negative SI (-4.8±0.9%), suggesting a small reduction in algal death during viral infection (**Fig. 3d**). Correspondingly, *A. stellipolaris* MB63 exhibited a high BI (48.9±2.7%), reflecting an almost 50% growth enhancement during viral infection (**Fig. 3g**), whereas *S. pontiacus* MB3 showed a growth deficit in the virus-infected condition (BI = -1.9±4.2%; **Fig. 3h**).

We extended this analysis to 10 bacteria isolated from the BAM consortium (**Table S1**; **Fig. S3-S4**). The three *Alteromonas* strains tested (MB63, MB67 and MB68) consistently exhibited strong effects on algal population decline (i.e., SI 12.0 - 16.6%; **Fig. 3i**) while simultaneously deriving substantial growth benefits from viral infection (i.e., BI 16.8 - 48.9%; **Fig. 3i**). We observed high variability in both metrics (i.e., SI -20.6 - 6.7% and BI -14.4 - 913.7%) for the other BAM-derived isolates (**Fig. S3-S4**).

By representing 19 *G. huxleyi*-associated bacterial isolates on a coordinate system defined by their SI and BI values (SI-BI phase space; **Fig. S3-S5**), we identified distinct bacterial ecological strategies within the algal bloom microbiome (**Fig. 3j**). We classified these bacterial taxa into four groups based on their position in this SI-BI phase space: (1) “Opportunistic pathogens” (high SI, high BI), exemplified by *Alteromonas* species, which both accelerate algal mortality and efficiently exploit the resulting metabolic niche; (2) “Consumers” (low SI, high BI), such as *Croceibacter atlanticus* MB69 and *Olleya* sp. MB6, which derive substantial growth benefits from viral infection without contributing significantly to algal mortality; (3) “Killers” (high SI, low BI), including *Roseovarius nubinhibens* MB34, which enhance algal mortality but gain no growth advantage; and (4) “Agnostic bacteria” (low SI, low BI), like *Marinobacter* sp. MB49, which neither increase algal mortality nor derive substantial benefit from viral infection.

### Synergistic bacteria harbor specific metabolic pathways

To identify genetic determinants of bacterial synergy in virus-induced algal demise, we sequenced and annotated the genomes of all bacterial isolates. We correlated the SI of each isolate with pathway completeness (the number of genes encoded in each KEGG pathway). We found significant positive correlations between SI and several metabolic pathways, most notably fatty acid degradation (Spearman’s ρ = 0.749, adj. p = 0.034), amino sugar and nucleotide sugar metabolism (ρ = 0.703, p = 0.044), branched-chain amino acid degradation (ρ = 0.760, p = 0.034), and trimethylamine metabolism genes belonging to the general methane metabolism pathway (ρ = 0.679, p = 0.047; **Fig. S6**; **Tables S2-S5**).

Detailed examination revealed that within the BAM consortium, the species with the highest SI values – all belonging to the genus *Alteromonas* — also encoded the complete enzymatic machinery for triacylglycerol (TAG) hydrolysis and subsequent fatty acid transmembranal transport and β-oxidation, including TAG lipase, *fadL*, *fadD*, *fadE*, *fadB*, *fadA*, and their homologs (**Fig. 4**). In contrast, species with low SI values (e.g., *Sulfitobacter pontiacus* MB3) showed several missing enzymes in the fatty acid degradation pathway. Within bloom-associated bacterial isolates, only *Alteromonas* and *Marinobacter* strains encode these metabolic genes (**Table S2**). While this may be a phylogenetic trait that is not causally related to SI or BI, these genera were recently shown to degrade algal lipids (*29*). Most importantly, *Alteromonas* encodes for a TAG lipase, an enzyme that releases free fatty acids from TAGs for cellular metabolism. This feature may be particularly advantageous during viral infection of *G. huxleyi*, as EhV infection remodels its lipid metabolism inducing TAG production and accumulation in intracellular lipid bodies (*30*). EhV infection further induces the release of extracellular vesicles that are enriched in TAGs (*31*), which can fuel bacteria that encode for TAG-specific catabolic enzymes.

**Figure 4.**
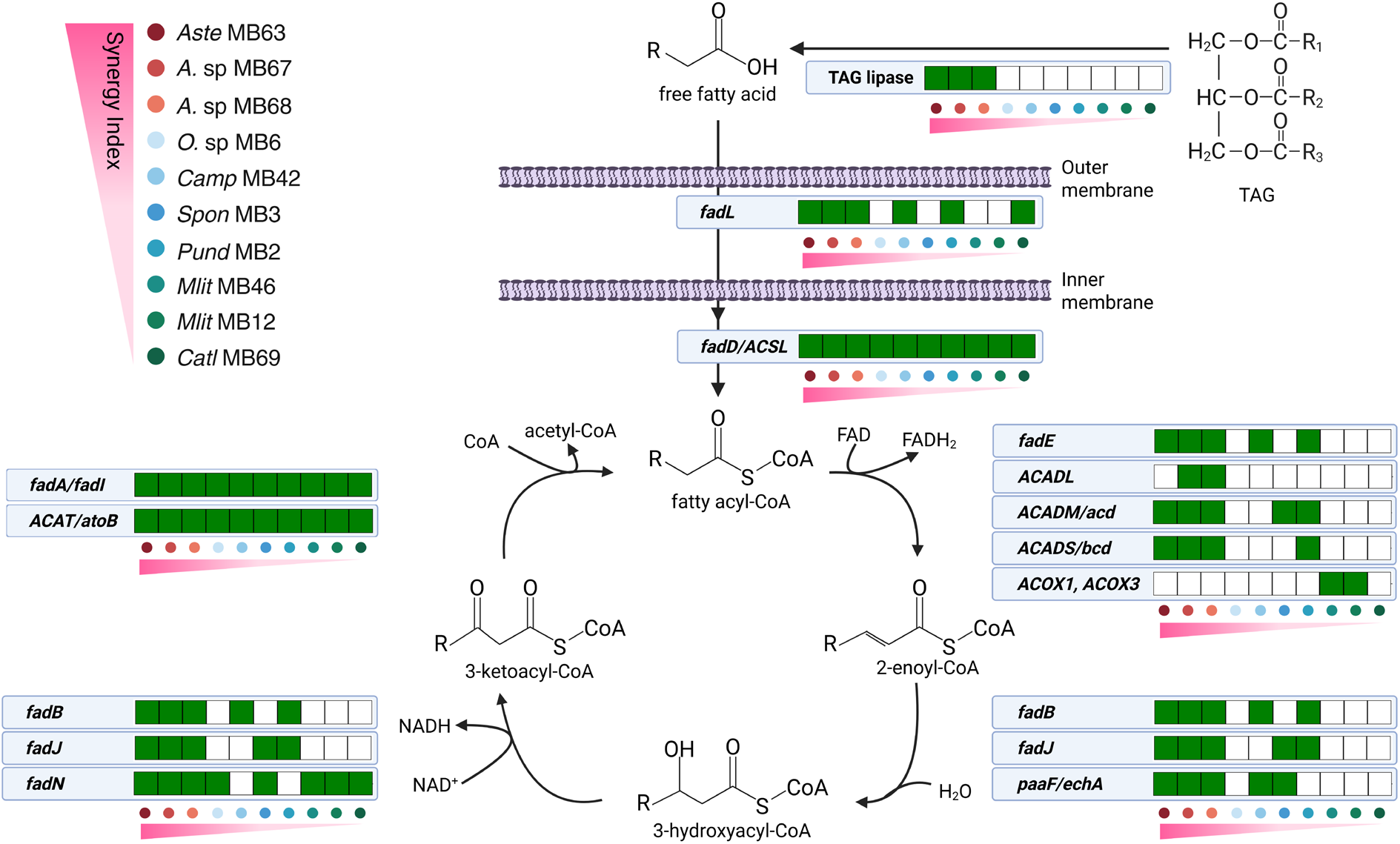
Fatty acid degradation pathway completeness correlates with bacterial synergy in algal mortality. KEGG functional analysis of ten isolated BAM members reveals enrichment of fatty acid degradation genes in synergistic bacteria (**Figures S6a**). The triacylglycerol (TAG) lipase that is not part of the pathway but provides its initial substrate is also included. Green boxes indicate the presence of genes, white indicates their absence (**Table S2**). The pink gradient represents SI values, with darker pink representing a higher synergistic effect.

### *Alteromonas* exhibit specialized ecological strategies during bloom demise

Having identified *Alteromonas* as key players in the viral-bacterial synergy during algal bloom demise, we investigated the ecological significance of this genus during natural bloom succession. To this end, we tracked algal, viral and bacterial dynamics in a natural environment, measured daily during *G. huxleyi* bloom succession in the mesocosm experiment (*18*). Following six days at low abundances (<5 × 10^5^ cells mL^-1^; *Pre-bloom* phase), *G. huxleyi* showed a steady increase in population (days 7-14; *Bloom* phase) and, upon viral infection (days 15-23; *Infection* phase), its population collapsed (**Fig. 5a**).

**Figure 5.**
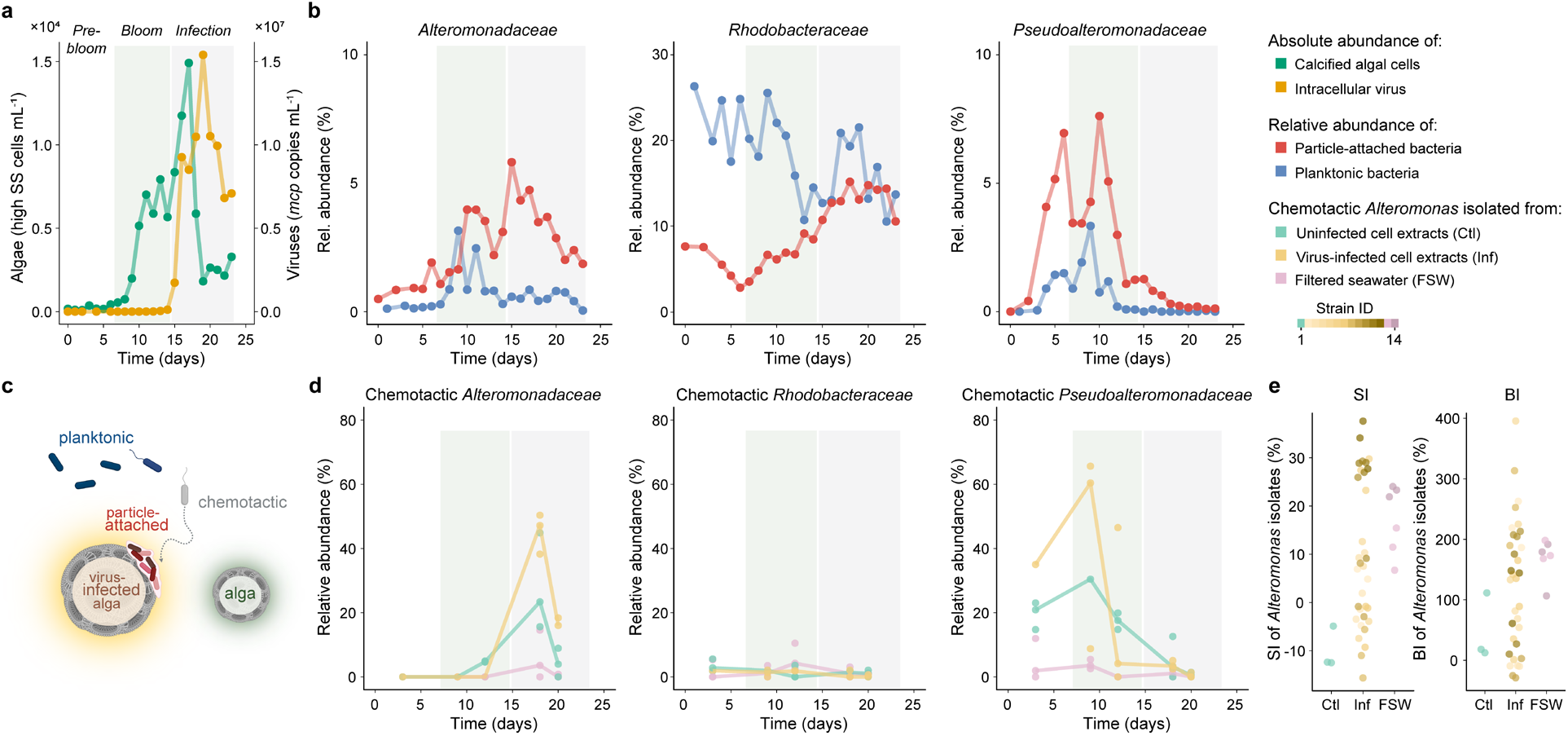
Members of the *Alteromonadaceae* family exhibit pronounced particle association and chemoattraction during virus-induced bloom demise. (a) Abundances of *G. huxleyi* (green) and intracellular EhV (yellow) in an induced mesocosm bloom defined a pre-bloom phase (days 1-6), a bloom phase (days 7-14) and infection phase (days 15-23). (b) 16S relative abundances of bacterial families (as named on the y-axis) in planktonic (<2 μm; blue) and particle-attached (>20μm; red) fractions. (c) Scheme of enrichment in *Alteromonadaceae* on particles during the infected bloom phase may be facilitated through chemoattraction towards virus-infected cells. (d) Relative abundance of bloom-associated *Alteromonadaceae*, *Rhodobacteraceae* and *Pseudoalteromonadaceae* in the chemotactic bacterial communities that entered the *in-situ* chemotaxis assays (ISCAs) deployed in the mesocosm during the *G. huxley*i bloom. Virus-infected (yellow) and uninfected (green) *G. huxleyi* cell extracts were used as chemoattractants in the ISCAs (n = 3). Mesocosm water was used as a reference (pink). (e) SI and BI of chemotactic *Alteromonas* strains isolated from ISCA wells containing uninfected *G. huxleyi* extracts (Ctl, one isolate) and virus-infected extracts (Inf, 11 isolates), compared to members of the BAM community (Bloom, two isolates; see Fig. 3i). Each strain was tested in biological triplicates. Complete SI and BI plots for these 14 different *Alteromonas* isolates are presented in **Figs. S10-S11**.

We used 16S rRNA amplicon sequencing to track bacterial abundances in two size fractions to distinguish between planktonic (0.2 - 2 μm) and particle-attached (20 - 200 μm) populations (**Fig. 5b**; **Fig. S7**). We observed significantly higher relative abundances of particle-attached *Alteromonadaceae* in all bloom phases (Kruskal-Wallis test; p < 0.05; **Fig. S8a**). During the onset of viral infection and subsequent bloom demise (days 15-23), *Alteromonadaceae* became on average 6-fold enriched in the particle-attached fraction compared to the uninfected bloom phase (Welch test; p < 0.05; **Fig. S8b**), suggesting *Alteromonadaceae* specifically colonize algal aggregates generated during infection. Viral infection of *G. huxleyi* blooms enhance the production of transparent exopolymer particles (TEP) up to 4.5 fold (*18*), generating a sticky glycan coating that catalyzes aggregate formation (*32*, *33*).

The dynamics of other bloom-associated bacterial families were markedly different from *Alteromonas* during the *G. huxleyi* bloom (**Fig. 5b**). *Rhodobacteraceae* showed up to 3-6x higher abundances in the planktonic compared to the particle-attached population during the pre-bloom and bloom phase (Kruskal-Wallis test; p < 0.05; **Fig. S8c**) with an increase in the particle-attached fraction throughout the phytoplankton bloom succession (**Fig. S8d**). *Pseudoalteromonadaceae* exhibited an increase in relative abundances in both planktonic and particle-attached fractions during the pre-bloom phase followed by a decrease during the bloom phase (**Fig. 5b**), suggesting that these bacteria are part of a different ecological niche.

To further investigate whether *Alteromonadaceae* actively seek out virus-infected cells during the bloom and demise of *G. huxleyi*, we deployed the *in-situ* chemotaxis assay (ISCA) (*34*) at multiple times during bloom succession. The ISCA is a microfluidic device containing an array of micro-wells that are filled with different chemicals. Here we loaded the ISCA wells with cell extracts from virus-infected (‘Inf’), or uninfected (‘Ctl’) laboratory *G. huxleyi* cultures, as well as filtered seawater (‘FSW’) acting as a reference. We then deployed the ISCAs in the mesocosm water, allowing the chemical extracts to diffuse out of the wells, therefore creating concentration gradients that can attract chemotactic bloom-associated bacteria into the wells. We characterised the composition of the chemotactic bacterial communities in the ISCA wells using 16S rRNA gene amplicon sequencing (**Fig. 5c, d**).

*Alteromonadaceae* were enriched in ISCA wells containing extracts from both ‘Inf’ (yellow line) and ‘Ctl’ (green line) *G. huxleyi* during the peak of viral infection in the demise phase of the bloom (days 18 and 20), accounting for up to nearly half (45.3±6.3%) of all chemotactic bacteria in the ‘Inf’ wells on day 18 (**Fig. 5d**). Notably, *Alteromonadaceae*, specifically *Alteromonas* (**Fig. S9**), were significantly enriched in the ISCA wells containing virus-infected *G. huxleyi* cell extract compared to the seawater reference (6.1±7.6%; pink line; Welch’s ANOVA, p < 0.01), suggesting they are attracted by the algal-derived metabolites. *Alteromonadaceae* were 19 times enriched (13.1±19.6%) in ‘Inf’ ISCA wells compared to their relative abundance in the mesocosm bloom (0.7±0.7%). This contrasts with other microbial families, including those whose abundance in the mesocosm bloom was higher. For example, *Rhodobacteraceae* accounted for 18.3±4.8% of planktonic bacteria, which is 26 times higher than the proportion of *Alteromonadaceae* during the overall bloom succession (**Fig. 5b**) but had a 20 times lower relative abundance in ‘Inf’ ISCA wells (0.9±1.0%; **Fig. 5d**). *Pseudoalteromonadaceae* were also enriched in ISCA wells containing *G. huxleyi* extracts but primarily during the pre-bloom and bloom phases (days 3, 9 and 12), accounting for up to 44.9±31.4%, rather than specifically during the viral infection phase at which *Pseudoalteromonadaceae* accounted for only 5.2±6.6% in ‘Ctl’ and 3.6±1.3% in ‘Inf’ ISCA wells (days 18 and 20; **Fig. 5d**), which is consistent with their higher relative abundances in bloom water during these early phases (**Fig. 5b**).

We sought to determine the synergy and benefit of strains that were selectively chemoattracted to *G. huxleyi*. To this end, we isolated 12 *Alteromonas* strains directly from the ISCA wells and quantified their SI and BI using our tripartite model system (**Table S1**; **Figs. S10**-**S12**). Intriguingly, some of the most synergistic and benefitting *Alteromonas* strains (highest SI and BI) were isolated from the ISCA wells containing the virus-infected extracts (**Fig. 5d**), suggesting that *Alteromonas* evolved specific traits that allow it to occupy the metabolic niche that is created by viral infections in *G. huxleyi* blooms.

## Discussion

The viral shunt is a key ecological process that recycles photosynthetic biomass generated in the sunlit surface ocean, providing nutrients and metabolites that fuel the marine microbiome. Our findings demonstrate that this process can selectively modulate the bloom’s microbiome composition and thereby expedite bloom demise. This viral-bacterial synergy helps to explain why entire *G. huxleyi* blooms collapse synchronously despite only 10-30% of the cells being actively infected by EhV (*24*, *25*). Viral infection may therefore be a trigger for additional pathogens expediting bloom demise.

Here, we introduced a new experimental framework for studying complex microbial interactions. By generating a synthetic community composed of alga, bacteria and viruses isolated from an algal bloom, we experimentally quantified their tripartite dynamics. To this end, we developed two novel quantitative indices: the Synergy Index (SI) measuring killing interaction strength, and the Benefit Index (BI) assessing bacterial fitness gain. We measured close to 20% increase in viral-induced algal mortality in the presence of the bloom-associated microbiome, compared to viral infection alone, and a nearly 400% increase in bacterial benefit.

This quantitative approach allowed us to disentangle these complex natural microbial communities in the lab and identify key players in algal bloom demise. We revealed *Alteromonas* bacteria as major drivers of this globally important process. This cosmopolitan genus was shown to exhibit remarkable lifestyle plasticity (*35*, *36*), including pathogenicity (*36*, *37*) and chemoattraction towards photosynthetic microbes (*38*). Using our quantitative framework, we show that these bacteria switch from commensals to opportunistic pathogens during viral infection of coccolithophore blooms, exhibiting high killing interaction strength (SI) and fitness gain (BI). In addition, *Alteromonas* displayed high chemoattraction towards the unique chemical seascape of infected cells and accumulated in sinking particles derived from bloom demise. Thus, *Alteromonas*, despite its low abundance in healthy algal-associated populations, is a keystone species uniquely adapted to viral-induced algal bloom demise.

We propose that the ephemeral nature of the release of unique and nutritious metabolites during virus-induced bloom demise, selectively recruits bacterial taxa with such specialized adaptive traits (as was shown in bipartite algal-bacterial interactions (*39*)). Viral infection of *G. huxleyi* generates unique metabolic seascapes, producing specific halogenated compounds and lipids (e.g., TAGs consumed by *Alteromonas*; **Fig. 4**) upon cell lysis (*15*, *30*, *40*, *41*). Viral infection also stimulates the production of transparent exopolymer particles (TEPs), which increase cell stickiness and facilitate bacterial adherence (*18*, *32*, *42*). We showed that *Alteromonas* is significantly enriched in particle-associated microbial communities during algal bloom demise.

We complemented the laboratory model for viral-bacterial synergy with an *in-situ* chemotaxis assay (ISCA), allowing us to quantify bacterial behaviour directly at different phases of bloom succession. We tailored this experiment specifically to detect, isolate and characterize bacteria that were attracted to virus-infected *G. huxleyi* cells. During bloom demise, *Alteromonas* demonstrated specific attraction towards the chemicals released by infected *G. huxleyi* cells (**Fig. 5d**), which likely led to the increase in particle association observed (**Fig. 5b**). Indeed, *Alteromonas* were previously shown to exploit adhesive algal surfaces for colonization (*43*). Furthermore, *Alteromonas* strains that were specifically attracted to infected algae also exhibited high SI and BI (**Fig. 5e**).

Our results demonstrate viral-bacterial synergy at the community level, raising intriguing questions regarding the specific cellular mechanisms that drive both the synergistic killing (SI) and the associated bacterial fitness benefits (BI). We propose four non-mutually exclusive scenarios that may underlie this phenomenon. First, viral infection may trigger the release of infochemicals that induce bacterial pathogenicity toward both infected and uninfected algal cells, thereby increasing mortality beyond the directly virus-infected population. Second, bacteria may accelerate the death of virus-infected algae by efficiently metabolizing essential cellular components, such as membrane lipids, thereby compromising cellular integrity. Third, bacterial metabolites or signaling molecules produced during viral infection may interfere with algal cell division, reducing population growth rates even in the absence of direct cell lysis. Fourth, bacteria that proliferate by exploiting viral lysate may competitively exclude algal cells for essential nutrients, creating resource limitation that reduces algal growth. Future mechanistic studies will be essential to distinguish between these scenarios and identify the precise molecular drivers of this ecologically significant tripartite interaction.

This is the first demonstration of how co-infection by two agents, namely viruses and bacteria, can act in synergy in the marine environment, akin to co-infections that are major mortality agents in human diseases (*27*, *28*). Bacterial pathogenicity has been shown in the past to depend on contextual environmental parameters such as temperature (*44*) and the growth phase of the algal host (*45*, *46*). Here, we introduce viral infection as a major environmental factor that may be the trigger of bacterial pathogenicity driving bloom demise. We propose that the biogeochemical implications of these tripartite interactions are profound. Viral-bacterial synergy (SI) expedites bloom demise, likely reducing the amount of carbon fixed during algal blooms. Bacterial growth due to algal bloom demise (BI) can affect the balance between the viral shunt (increase the recycling of carbon in the ocean surface, reflected in a high BI) and the viral shuttle (exporting carbon to ocean depths via sinking particles). Deciphering the nature of algae-virus-bacteria interactions is critical given the role of large-scale oceanic blooms in carbon and sulfur cycling, as well as climate regulation.

## Materials and methods

### Isolation of alga, virus and BAM from a *G. huxleyi* bloom

*G. huxleyi* strain RCC6946, the virus EhV-M1, and the bloom-associated microbiome (BAM) were isolated from an induced *G. huxleyi* bloom during the AQUACOSM VIMS-Ehux mesocosm experiment conducted near Bergen, Norway, in 2018 (*18*). For algal isolation, a water sample collected from mesocosm bag 4 during the bloom phase (day 13, 06.06.2018) was used for single-cell isolation within two weeks of sample collection at the Roscoff Culture Collection (RCC). The bloom-associated microbiome (BAM) was isolated from mesocosm bag 7, which exhibited the highest increase in viral abundance during *G. huxleyi* bloom demise. A water sample was collected during the demise phase (day 23, 16.06.2018), filtered through GF/F glass microfiber filters (∼0.7 µm nominal pore size), and the filtrate cryopreserved in 15% glycerol (final concentration). The lytic virus EhV-M1 was isolated from the same mesocosm experiment using *G. huxleyi* CCMP 374 as susceptible host, and its genome was sequenced (*47*).

### Maintenance of *G. huxleyi* cultures and alga-virus infection experiments

The *G. huxleyi* strain RCC6946 was maintained in batch cultures at 18°C with a 16:8-h light:dark cycle. Light was provided at 100 μmol photons m^-2^ s^-1^ using cool white LEDs. The medium was composed of autoclaved filtered seawater (FSW) supplemented with modified K/2 medium (*48*), modified as described in **Table S6**). Weekly dilutions were performed at a 1:10 ratio with fresh medium to maintain the cells in exponential growth. In parallel to the xenic culture, an axenic culture was derived and maintained by adding the antibiotics ampicillin and kanamycin at 100 μg mL^-1^ and 50 μg mL^-1^ final concentration, respectively. The absence of bacteria in axenic cultures was verified periodically by plating 100 μL on Difco Marine Agar 2216 plates (MA; BD Diagnostics) and enumerating bacteria using flow cytometry.

For viral infections, exponentially growing algal cultures at about 8 × 10^5^ cells mL^-1^ were infected with EhV-M1 (freshly propagated on RCC6946 and used within 7 days) at a multiplicity of particles (MOP) of 1:2 of alga:virus. For testing the effect of antibiotics on algal mortality, ampicillin and kanamycin were added to xenic RCC6946 cultures at 100 μg mL^-1^ and 50 μg mL^-1^ final concentration, respectively, simultaneously with viral co-infection at a ratio of 1:5 of algal cells to viral particles. For experiments with the BAM or bacterial isolates, the antibiotics concentration in the algal cultures was diluted at least 10,000-fold prior viral infection by dilution into fresh medium without antibiotics.

### Enrichment of the BAM on *G. huxleyi*

To enrich the BAM on *G. huxleyi*, the cryopreserved BAM was grown on diverse media containing organic material derived through exudation, mechanical lysis and viral lysis of algal cells, as detailed in the following. To collect Exponential Conditioned Media (Exp. CM) and Sonication Lysates (SL), algal cultures were grown to their exponential phase at about 8 × 10^5^ cells mL^-1^ and centrifuged at 600×g for 5 min. The supernatant was filtered through 0.22 μm PES filters and used as ‘Exp. CM’. The cell pellet was physically disrupted through probe sonication (5 cycles of 30 sec followed by 30 sec rest). Once sonicated, the lysate was resuspended in 500 mL K/2 medium, filtered through 0.22 μm PES filters and used as ‘SL’. To collect Viral Free Lysate (VFL), algal cultures were infected with EhV-M1 as described above. Four days post-infection, the lysed cultures were filtered sequentially through 2 μm cellulose membrane filters, 0.22 μm PES filters and 100 kDa centrifugal filters, and used as ‘VFL’. All three filtrates were kept at 4°C until further use within 24 hours. The BAM glycerol stock was directly inoculated into 30 mL of each filtrate in triplicates and incubated for 3 days in an orbital shaker at 18°C. Triplicates were combined and aliquots of 0.5 mL were cryopreserved in 15% glycerol (final concentration).

### Tripartite algae-virus-bacteria interaction experiments

Algal cultures were infected with EhV-M1, as described above, and co-inoculated with an overnight culture of BAM. BAM was inoculated from a glycerol stock into Difco Marine Broth 2216 (MB; BD Diagnostics) and incubated overnight at 28°C and 168 RPM. The overnight culture was washed three times with 0.2 µm-filtered FSW by centrifugation (10,000×g, 1 min), enumerated using flow cytometry, and inoculated at 1 × 10⁴ bacterial cells mL⁻¹ into the virus-infected algal cultures. All experiments were performed in biological triplicates, and statistical significance was assessed using a two-way ANOVA followed by Sidak’s multiple comparisons test (p < 0.05).

### Enumeration of algae, bacteria and viruses by flow cytometry

Flow cytometry analyses were performed using a CytoFLEX S flow cytometer (Beckman Coulter). Algal cells were identified as high forward scatter (FSC), high chlorophyll population by plotting FSC (height threshold = 4 × 10^4^ arbitrary units [A.U.]) versus chlorophyll fluorescence (ex.: 561 nm, em.: 665-715 nm). Only cells with a fluorescence >4 × 10^4^ A.U. were enumerated as live cells (**Fig. S13a-b**). For bacteria and virus counts, samples were fixed with glutaraldehyde (0.5% final concentration) for at least 30 min at 4°C, then snap-frozen in liquid nitrogen and stored at -80°C until analysis. After thawing, samples were stained with SYBR Gold (Invitrogen; diluted 1:10^4^ in Tris-EDTA buffer) for 20 min at 80°C and analysed by flow cytometry (ex.: 488 nm, em.: 500-550 nm) by plotting Violet SSC (area threshold = 1 × 10^3^ A.U.) versus FITC (area threshold = 1 × 10^3^ A.U.; **Fig. S13c-d**). All flow cytometry data were processed in CytExpert 2.4 (Beckman Coulter), and statistical analyses were conducted in GraphPad Prism (v10) and R (v4.2.0).

### Bacterial community analysis using 16S rRNA amplicon sequencing

Tripartite algae-virus-bacteria interaction experiments were set up in five replicates at 400 mL. Aliquots of 40 mL were filtered by gravity through autoclaved GF/C filters (47 mm; Whatman), and the filtrate collected on autoclaved 0.22 µm PVDF filters (47 mm; Durapore, Mercury Millipore). The filters were directly transferred to the lysis buffer of DNeasy PowerWater Kit (Qiagen), frozen in liquid nitrogen, and stored at −20°C until further processing. DNA extraction was performed according to the manufacturer’s instructions. The 16S rRNA gene was amplified using primers targeting the V4 region, 515F (49); GTGYCAGCMGCCGCGGTAA) and 806R (50); GGACTACNVGGGTWTCTAAT), under the following PCR conditions: initial denaturation at 94°C for 3 min, followed by 35 cycles of denaturation at 94°C for 45 sec, annealing at 50°C for 1 min, and elongation at 72°C for 1 min and 30 sec. A final elongation step was performed at 72°C for 10 min to ensure complete extension of the amplified products. Sequencing was performed using the Illumina MiSeq platform at the RUSH Genomics and Microbiome Core Facility.

The 16S rRNA amplicon sequencing data were processed using the DADA2 pipeline (v1.28.0; (*51*) in R (v4.3.2) to perform quality filtering, error correction, and amplicon sequence variant (ASV) inference. Raw sequencing reads were quality-checked using the plotQualityProfile function, and low-quality bases were trimmed using the filterAndTrim function with truncation parameters set based on quality score distributions. Error rates were learned from the dataset using the learnErrors function, and sequences were dereplicated using derepFastq. ASVs were inferred using the dada function, followed by chimera removal with removeBimeraDenovo. Taxonomic classification was performed using the SILVA_nr99 reference database (v138.1; (*52*), and ASV tables were generated for downstream analyses. The average sequencing depth per sample was 67,413±26,969 (min 24,150 and max 132,743 reads). Chloroplast reads accounting for 28±23% (max 73%) were removed. The processed data were then analyzed using the Phyloseq package (v1.44.0; (*53*) in R to visualize microbial community composition, calculate alpha and beta diversity metrics, and perform statistical comparisons of microbial communities across experimental conditions.

### Calculation of Synergy Index (SI) and Benefit Index (BI)

SI was determined by comparing the area under the growth curve (AUC) of virus-infected *G. huxleyi* in the presence and absence of bacteria (the entire growth curves were used). The AUC was calculated using the ‘trapz’ function of the ‘NumPy’ package in Python. We chose to focus on AUC because of four main reasons: First, It is biologically meaningful and captures the cumulative biomass change that would occur in natural bloom systems, reflecting the total “ecological impact”. For example, the total amount of carbon fixed. Second, it captures complete temporal dynamics rather than relying on a single timepoint or summary statistic that would miss critical dynamics. Third, it is more robust against experimental variability, as it takes into account 3 measurements from each of the experiment’s days. Finally, the normalization of AUC difference by another AUC, creates a dimensionless metric that allows direct comparison across different bacterial strains and experimental conditions even if their baseline levels are different. The SI was calculated as follows:

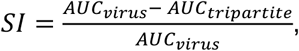

where:

- *AUC_virus_* represents the algal population dynamics under viral infection alone.
- *AUC_tripartitte_* represents the algal population dynamics under co-infection with both viruses and bacteria.

BI quantifies bacterial growth differences over time by comparing the AUC in tripartite conditions versus bacterial growth alone. BI is calculated as:

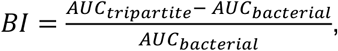

where:

- *AUC_tripartitte_* represents the bacterial growth over time when co-cultured with virus-infected *G. huxleyi*.
- *AUC_bacterial_* represents bacterial growth over time in the presence of *G. huxleyi* but without viral infection.

Synergy Index (SI) uncertainty was quantified using analytical error propagation with n=3 replicates per condition. For each condition, mean area under curve (AUC) and standard error (SE = σ/√n) were calculated. SI variance was determined using the delta method for the ratio SI = (A - B)/A, where A = μ_control and B = μ_infected: Var(SI) = (B/A^2^)^2^ × Var(A) + (1/A)^2^ × Var(B). Standard error of SI was calculated as SE_SI = √Var(SI), and 95% confidence intervals were derived using the t-distribution with degrees of freedom = n_control + n_infected - 2. Results are reported as SI ± SE.

### Isolation of bacteria from BAM and induced *G. huxleyi* blooms

For bacterial isolation, glycerol stocks were sampled at different days of the induced *G. huxleyi* blooms and following various enrichments (**Table S1**). Briefly, filtered seawater samples (using GF/F and 10 µm filters) were collected during the bloom (day 13) and demise (day 21 and day 23) phases and cryo-preserved as glycerol stocks at 15% final concentration. The BAM glycerol stock was prepared as described above. To enrich alga- and viral infection-associated taxa, the mesocosm and BAM glycerol stocks were inoculated into uninfected and virus-infected cultures of *G. huxleyi* before isolation. To enrich G. *huxleyi*-associated taxa, xenic cultures of RCC6946, which contain bacteria that were co-isolated with the algal cell from the mesocosm bloom, were infected with EhV-M1 before isolation. From overnight cultures of the glycerol stocks or from enrichment cultures, 100 µL were plated at 100×, 1,000×, and 10,000× dilution on MA and incubated at 28°C, allowing for single colony isolation. Selected colonies were streaked onto fresh MA plates three times or until a uniform colony morphology was observed, ensuring the purity of the bacterial isolate. Colonies were inoculated in 3 mL MB medium and grown overnight at 28°C with shaking at 160 RPM. Glycerol stocks were prepared at 15% final concentration, snap-frozen in liquid nitrogen and stored at -80°C.

For sequencing the complete 16S rRNA gene, cells from pure colonies were lysed using a buffer containing TE and 0.1% Triton-X100, followed by heating at 95°C for 5 minutes. Cell lysates were centrifuged at 13,300 RPM for 10 min, and 2 µL of the supernatant was used as a DNA template for PCR amplification. The 16S rRNA gene was amplified using the universal bacterial primers 16S-27F (AGAGTTTGATCMTGGCTCAG) (*54*) and 16S-1492R (CGGTTACCTTGTTACGACTT) (*55*). The PCR reaction was carried out in 20 µL using 2× HY-Taq (HyLabs) or REDTaq (Sigma) Ready Mix. PCR amplification was performed using the following conditions: initial denaturation at 94°C for 3 min, followed by 25–35 cycles of denaturation at 94°C for 30 sec, annealing at 57°C for 30 sec, and elongation at 72°C. A final elongation step was conducted at 72°C for 10 min, followed by a hold at 12°C. PCR products were visualized in 1% w/v agarose gel with ethidium bromide. Amplicons were purified with the ExoProStar 1-Step enzymatic cleanup kit using 5 µL aliquot of the PCR reaction and incubated at 37°C for 15 min, followed by enzyme inactivation at 80°C for 15 min. For Sanger sequencing, 1-2 µL of the purified amplicons were analyzed by the Life Sciences Core Facilities at the Weizmann Institute of Science. All 16S rRNA gene sequences were classified at the genus level using the SINA classifier tool (v1.2.12) within the SILVA rRNA database (*56*); **Table S1**).

### Genomic sequencing of bacterial isolates and metabolic pathway analysis

The genome of *Sulfitobacter* sp. D7 was sequenced as described previously (*57*). All bacterial isolates from the mesocosm (**Table S1**) were inoculated from glycerol stocks in MB medium and grown overnight at 28°C with agitation (160 RPM). Cultures were centrifuged at 12,000 RPM at room temperature for 1 min, the cell pellet washed 3× with FSW, and frozen in liquid nitrogen. Genomic DNA was extracted following a modified version (*58*) of a previously described protocol (*59*). High molecular weight of the genomic DNA was verified by gel electrophoresis or Genomic DNA ScreenTape Analysis using TapeStation 4150 (Agilent Technologies), while DNA concentration and purity were validated using a Qubit fluorometer with the dsDNA HS Assay Kit (Thermo Fisher Scientific) and NanoDrop ND-1000 spectrophotometer (Thermo Fisher Scientific), respectively. If necessary, samples were concentrated using the NEBNext Ultra II FS DNA Library Prep Kit (New England Biolabs) or AMPure XP beads (Beckman Coulter).

For Illumina short-read sequencing, shotgun metagenome libraries were prepared using the NEBNext Ultra II FS DNA Library Prep Kit for Illumina (New England Biolabs) with enzymatic shearing, according to the manufacturer’s instructions. The final size-selected pool was sequenced in an Illumina NovaSeq6000 instrument with an S2 flow cell, employing 2×150 base reads. Short reads were quality trimmed using Trimmomatic (v0.39) with the following parameters: ILLUMINACLIP:TruSeq3-PE-2.fa:2:30:10, LEADING:10, TRAILING:10, SLIDINGWINDOW:4:15, MINLEN:50. Reads that passed quality trimming were used for assembly construction with SPAdes (v3.13.0), using default k-mer sizes (21 bp, 33 bp, 55 bp, and 77 bp). Contigs were filtered based on mean coverage (≥3x) and length (≥300 bp) and clustered into bins using MaxBin2 (v2.2.7) with the default parameters. Assembly quality was assessed using CheckM2 (v1.0.2) with default parameters. Multiple bins of the same isolate were concatenated into single genomes if they complemented one another based on completeness and had similar taxonomic annotation. Genes were predicted from the genomes using Prodigal (v2.6.3) in metagenomic mode. Predicted genes were filtered based on length (≥300 bp) and the presence of a start codon at the first trimer, and the absence of a premature stop codon. Genes that passed filtration were clustered using MMseqs2 (v13.45111) with the following parameters: coverage mode 0, minimum coverage 0.8, and minimum sequence identity 0.9. Representative genes were annotated using eggNOG-mapper (v2.1.3) and used for downstream analysis.

For PacBio long-read sequencing, multiplexed microbial libraries were prepared, using SMRTBell Express Prep Kit 2.0 (Pacific Biosciences). A single PacBio sequencing library was constructed that was used for SMRTbell template preparation as described in the protocol (Pacific Biosciences). The resulting SMRTbell template was annealed to the sequencing polymerase, using a Sequel binding kit 2.0. Sequencing was conducted with a 12pM on-plate concentration using a Sequel I System in CLR mode and a single SMRT Cell (1M; PacBio). PacBio raw reads (polymerase reads) were processed using the SMRT Link software (v8.0). Genomes were assembled using Trycycler v0.5.5 and Flye v2.9.6 with default parameters. Assembly quality was assessed using CheckM2 (v1.0.2) with default parameters. Genes were predicted, filtered, clustered and representative genes annotated as described above.

Nanopore sequencing was performed by Plasmidsaurus following their standard bacterial genome sequencing service using the Oxford Nanopore Technology with custom analysis and annotation.

All assembled genome sequences were classified at the species level using the GTDB-Tk software toolkit v2.4.1 with GTDB database version 220 (*60*); **Table S1**).

To assess the relationship between bacterial metabolic pathways and bacterial synergy across all isolated strains from BAM and the *G. huxleyi* bloom (n = 19), all KEGG pathways and their constituent KEGG Orthologs (KOs) were retrieved via KEGG API. Using the eggNOG-based genome annotations for each taxon, pathway coverage (%) was calculated as the proportion of KOs present in a strain relative to the total KOs in that pathway. Pathway coverage was then correlated with the synergy index using Spearman’s rank correlation, and the resulting p-values were adjusted for multiple testing with the Benjamini-Hochberg false-discovery rate procedure.

### Enumeration of algae and viruses in *G. huxleyi* blooms

The abundances of *G. huxleyi* and intracellular EhV throughout the succession of the induced *G. huxleyi* bloom in mesocosm bag 7 were enumerated as described previously (*18*). Briefly, for the enumeration of *G. huxleyi* cells, water samples were pre-filtered using 40 µm cell strainers and immediately analyzed with an Eclipse iCyt flow cytometer (Sony Biotechnology). Calcified *G. huxleyi* cells were identified as a population of cells with high side scatter and high chlorophyll (ex: 488 nm, em: 663-737 nm). For the enumeration of intracellular EhV, water samples were pre-filtered through 20 µm and DNA was extracted from 2 µm filters. EhV abundance was determined by qPCR using the major capsid protein (*mcp*) gene. Absolute abundances were calculated using a calibration curve of DNA at known concentrations.

### 16S rRNA amplicon sequencing of planktonic and particle-attached bacteria

Planktonic (>0.2 µm and <2 µm) and particle-attached (>20 µm- and <200 µm) fractions of the bacterial community were assessed throughout the succession of the induced *G. huxleyi* bloom in mesocosm bag 7 using 16S rRNA amplicon sequencing, as described previously (*18*). In brief, water samples were pre-filtered through 200 µm, sequentially filtered through 20 µm, 2 µm and 0.22 µm polycarbonate filters (Isopore, 47 mm; Merck Millipore), and immediately frozen in liquid nitrogen. DNA extraction of the 20 µm and 0.22 µm filters was performed with the DNeasy PowerWater kit (Qiagen), and sequenced following the EMP 16S amplicon protocol using the 515F-806R primers (*61*) including degeneracy to reduce bias towards Thaumarchaeota and SAR11 (*49*, *50*). Data analysis for planktonic bacteria was performed using the DADA2 pipeline and ASVs annotated with the RDP classifier. The average sequencing depth per sample and chloroplast read proportion for the planktonic community was reported previously (*18*). Data analysis for particle-attached bacteria was performed using the DADA2 pipeline and ASVs annotated with the SILVA_nr99 reference database (v138.1). The average sequencing depth for the particle-associated size fraction in bag 7 was 20,734±6,517 (min 8,057 and max 29,629) reads per sample. Chloroplast reads accounting for 7±8% (min 1% and max 36%) in the particle-attached fraction were removed. Pairwise Welch t-tests were performed for the log_2_ fold change (FC) of particle-associated to planktonic bacterial fractions between the pre-bloom (days 0-6), bloom (days 7-14) and viral infection phase (days 15-23).

### Chemotaxis assay and 16S rRNA amplicon sequencing of chemotactic bacteria

To prepare cell extracts for *in-situ* chemotaxis assays (ISCA), *G. huxleyi* CCMP 2090 was grown in 19 L modified K/2 medium (**Table S6**) in two carboys with stirring and aeration. One exponentially growing culture (1.2 × 10^6^ cells mL^-1^) was infected with EhV 201 at a MOP of 1:1 while the other culture served as uninfected control. The viruses were propagated on CCMP 2090, washed and concentrated on 100 kDa Amicon-ultra filters (EMD Millipore), and stored at 4°C until use within two months. Cells were counted daily using a Multisizer 4 Coulter Counter (Beckman Coulter). At 48 hours post-infection, the uninfected (4.4 × 10^6^ cells mL^-1^) and infected (0.9 × 10^6^ cells mL^-1^) cultures were filtered onto pre-combusted GF/C filters (47 mm; Whatman) using gentle vacuum (600 mbar under-pressure). Filters were immediately transferred to 500 mL pre-cooled (-20°C) methanol (HPLC grade, J.T. Baker) and extracted overnight at 4°C while shaking. Extracts were sonicated 30 min and filtered through pre-combusted GF/C filters before being evaporated to 20 mL using a Rotovap (30°C, 70-100 RPM). Extracts were centrifuged at 20,800×g for 10 min at 4°C and the supernatants evaporated to complete dryness using a speedvac (28°C, 1,980 RPM). Dried extracts were re-dissolved in MeOH to reduce the salt load, aliquoted to eight centrifuge tubes, dried using a speedvac, and stored at -80°C until further use. The dry weight was 57.5±0.3 mg and 38.3±0.5 mg for the uninfected and infected extracts, respectively.

ISCAs containing the *G. huxleyi* cell extracts were deployed five times throughout the mesocosm bloom succession, namely at days 3, 9, 12, 18 and 20. For each deployment, 12.5 mL of bloom water were collected per mesocosm bag (1-4; 50 mL in total), mixed and subjected to a quadruple-filtration process. Specifically, the mixed bloom water was first filtered through a 0.2 μm Millex FG (Millipore) to remove large particles, twice through a 0.2 μm Sterivex filter (Millipore), and finally through a 0.02 μm Anatop filter (Whatman). This quadruple filtration aimed to remove all microorganisms from the seawater. We confirmed the efficiency of the filtration procedure using flow cytometry and DNA sequencing (*62*). The filtered bloom water was used as a control in the ISCA, and was also used to resuspend the dried *G. huxleyi* extracts at a final concentration of 1 mg mL^-1^. The resuspended extracts were filtered through a 0.2 μm Millex FG (Millipore) to remove undissolved particles.

The working concentration of *G. huxleyi* extracts (i.e., 1 mg mL^-1^ of dry weight) is representative of environmental hotspots. Indeed, chemical concentrations decay exponentially with distance away from the ISCA port (*34*). This means that concentrations sensed by bacteria in the surrounding bloom water will be substantially lower than the one in the ISCA wells and within the same order of magnitude as the concentrations arising from the lysis of a phytoplankton cell (*63*).

Treatments (filtered bloom water and the two *G. huxleyi* cell extracts) were randomly allocated to an ISCA row (consisting of 5 wells). All wells in a row contained the same treatment, and each treatment was replicated on three different ISCAs, which were deployed in parallel to act as biological replicates (n = 3). Each ISCA was placed into a flow-damping acrylic enclosure, following a pre-established protocol (*64*). Each enclosure was filled with water coming from the first four mesocosm bags (i.e., bags 1-4), previously mixed in equal volume. The flow-damping enclosures were filled slowly with water from the inlet in the lower surface, and were sealed with a plug once full. The enclosures were then incubated *in-situ* at 1 m depth for 2 h (*62*). Following deployments, enclosures were retrieved and slowly drained. The volume of each ISCA well was retrieved using a sterile 1 mL syringe and 27G needle.

ISCA samples for DNA extraction were snap frozen immediately. DNA extraction was performed using a microvolume extraction approach (physical lysis method; (*65*). Libraries for 16S rRNA gene sequencing were generated for all samples using the Nextera XT DNA Sample Preparation Kit (Illumina) following a previously described protocol for low-input DNA libraries (*66*). Primers targeting the V4-V5 regions of the 16S rRNA gene were used, namely 515F (GTGYCAGCMGCCGCGGTAA) and 926R (CCGYCAATTYMTTTRAGTTT). Samples were then sequenced on an Illumina MiSeq platform. Libraries were pooled on an indexed sequencing run and yielded ∼400 bp per sample. To characterize the composition of bacterial communities, reads were first trimmed with Seqtk and pair-end were merged with FLASH (*67*). Adaptors were removed using CutAdapt (*68*). Denoising, dereplicating and taxonomy assignment were performed using DADA2 (*51*).

### Isolation of chemotactic *Alteromonas* for tripartite interaction experiments

To isolate chemotactic *Alteromonas* throughout the succession of the induced *G. huxleyi* bloom, we prepared agar plates made with 0.22-μm-filtered environmental bloom water and amended with 1 mg mL^−1^ of the corresponding *G. huxleyi* cell extracts. The agar plates were inoculated with the chemotactic bacterial community from the ISCA wells containing uninfected and virus-infected *G. huxleyi* extracts and incubated at fjord water temperature (10°C) until colonies appeared on the surface. Individual colonies were then grown in 10% MB in autoclaved seawater overnight at room temperature and at 10°C. Each isolate went through two additional rounds of purification via successive growth on plate and liquid cultures. Isolates were identified by 16S rRNA gene Sanger sequencing (*62*).

All *Alteromonas* strains (**Table S1**) were further used for genome sequencing and characterization of their SI and BI indices, as described above with slight modifications. In brief, overnight cultures were inoculated in MB medium at 0.1 OD_600nm_. Exponentially growing cultures at 1.0 OD_600nm_ were centrifuged and cell pellets of 15-30 mg re-dissolved in 0.5 mL Zymo DNA/RNA Shield. Genomic DNA extraction and Nanopore sequencing were performed by Plasmidsaurus following their standard bacterial genome sequencing service. Assembled genome sequences were classified at the species level using the GTDB-Tk software toolkit. Tripartite algae-virus-bacteria interaction experiments were performed and their SI and BI indices calculated, as described above. Day 6 was omitted due to technical reasons and thus SI and BI computed until day 5. Indices computed until day 5 and day 6 were correlated across strains and experiments (ρ_SI_ = 0.997; ρ_BI_ = 0.952).

## Acknowledgments

We thank all team members of the AQUACOSM VIMS-Ehux project for conducting the mescosom experiment, especially Jorun Egge, Aud Larsen, and Tatiana Tsagaraki. We thank Ian Probert and Martin Gachenot for isolating the algal strain RCC6946 during the mesocosm experiment. We are further grateful to Ari Isbi, Shai Duchin-Rapp and Alla Usyskin-Tonne for their help in Illumina sequencing of the isolated bacteria. The PacBio sequencing of selected bacterial genomes was performed by the WIS Genomics Unit. We thank Shaul Pollak and Uria Alcolombri, and the Vardi and Zeevi labs for their constructive comments.

## Author contributions

Conceptualization: CK, NBG, DS, DZ, AV

Investigation: ES and TSNZ performed tripartite interaction experiments. ES performed the 16S rRNA analysis for the tripartite interaction with BAM. NBG collected bacterial microbiomes from the mesocosm. IN, CK and TSNZ isolated bacterial strains. CK, TSNZ and DS performed DNA extractions for bacterial genome sequencing. ES and GT assembled bacterial genomes. ES performed the functional pathway analysis. ES and CK analyzed the 16S rRNA data from the mesocosm. CK, GS, EC and JBR performed ISCA experiments during the mesocosm. EC performed the 16S rRNA analysis for the ISCA experiments and isolated chemotactic bacteria.

Supervision: DZ, AV

Writing – original draft: ES, CK, DZ, AV

Writing – review: ES, CK, JBR, NBG, GT, GS, EC, RS, DS, DZ, AV

## Data and materials availability

16S rRNA sequences of the tripartite infection with BAM (Biosample SAMN49823070) and genome sequences of BAM-derived bacteria (accession numbers SAMN50000114-SAMN50000135 and SAMN50033443-SAMN50033450) are deposited under NCBI Bioproject PRJNA1287515. Flow cytometry data of the mesocosm *G. huxleyi* bloom has been deposited in Dryad (*69*). 16S rRNA sequences of the mesocosm have been deposited under NCBI Bioproject PRJNA694552.

## References

1. C. B. Field, M. J. Behrenfeld, J. T. Randerson, P. Falkowski, Primary production of the biosphere: integrating terrestrial and oceanic components. Science 281, 237–240 (1998).

2. P. G. Falkowski, R. T. Barber, V. Smetacek V., Biogeochemical controls and feedbacks on ocean primary production. Science 281, 200–207 (1998).

3. Azam, T. Fenchel, J. G. Field, J. S. Gray, L. A. Meyer-Reil, F. Thingstad, The ecological role of water-column microbes in the sea. Mar. Ecol. Prog. Ser. 10, 257–263 (1983).

4. M. J. Behrenfeld, R. T. O’Malley, D. A. Siegel, C. R. McClain, J. L. Sarmiento, G. C. Feldman, A. J. Milligan, P. G. Falkowski, R. M. Letelier, E. S. Boss, Climate-driven trends in contemporary ocean productivity. Nature 444, 752–755 (2006).

5. M. J. Behrenfeld, E. S. Boss, Resurrecting the ecological underpinnings of ocean plankton blooms. Ann. Rev. Mar. Sci. 6, 167–194 (2014).

6. Azam, Malfatti, Microbial structuring of marine ecosystems. Nat. Rev. Microbiol. 5, 782–791 (2007).

7. J. R. Seymour, S. A. Amin, J.-B. Raina, R. Stocker, Zooming in on the phycosphere: the ecological interface for phytoplankton-bacteria relationships. Nat. Microbiol. 2, 17065 (2017).

8. C. Kuhlisch, A. Shemi, N. Barak-Gavish, D. Schatz, A. Vardi, Algal blooms in the ocean: hot spots for chemically mediated microbial interactions. Nat. Rev. Microbiol. 22, 138–154 (2024).

9. C. A. Suttle, Marine viruses - major players in the global ecosystem. Nat. Rev. Microbiol. 5, 801–812 (2007).

10. C. P. D. Brussaard, Viral control of phytoplankton populations - a review. J. Eukaryot. Microbiol. 51, 125–138 (2004).

11. A. E. Zimmerman, C. Howard-Varona, D. M. Needham, S. G. John, A. Z. Worden, M. B. Sullivan, J. R. Waldbauer, M. L. Coleman, Metabolic and biogeochemical consequences of viral infection in aquatic ecosystems. Nat. Rev. Microbiol. 18, 21–34 (2020).

12. S. W. Wilhelm, C. A. Suttle, Viruses and nutrient cycles in the sea. Bioscience 49, 781–788 (1999).

13. M. B. Sullivan, J. S. Weitz, S. Wilhelm, Viral ecology comes of age. Environ. Microbiol. Rep. 9, 33–35 (2017).

14. M. G. Weinbauer, Ecology of prokaryotic viruses. FEMS Microbiol. Rev. 28, 127–181 (2004).

15. C. Kuhlisch, G. Schleyer, N. Shahaf, F. Vincent, D. Schatz, A. Vardi, Viral infection of algal blooms leaves a unique metabolic footprint on the dissolved organic matter in the ocean. Sci. Adv. 7, eabf4680 (2021).

16. Z. Zhao, M. Gonsior, P. Schmitt-Kopplin, Y. Zhan, R. Zhang, N. Jiao, F. Chen, Microbial transformation of virus-induced dissolved organic matter from picocyanobacteria: coupling of bacterial diversity and DOM chemodiversity. ISME J. 13, 2551–2565 (2019).

17. X. Ma, M. L. Coleman, J. R. Waldbauer, Distinct molecular signatures in dissolved organic matter produced by viral lysis of marine cyanobacteria. Environ. Microbiol. 20, 3001–3011 (2018).

18. F. Vincent, M. Gralka, G. Schleyer, D. Schatz, M. Cabrera-Brufau, C. Kuhlisch, A. Sichert, S. Vidal-Melgosa, K. Mayers, N. Barak-Gavish, J. M. Flores, M. Masdeu-Navarro, J. K. Egge, A. Larsen, J.-H. Hehemann, C. Marrasé, R. Simó, O. X. Cordero, A. Vardi, Viral infection switches the balance between bacterial and eukaryotic recyclers of organic matter during coccolithophore blooms. Nat. Commun. 14, 510 (2023).

19. M. A. Moran, E. B. Kujawinski, W. F. Schroer, S. A. Amin, N. R. Bates, E. M. Bertrand, R. Braakman, C. T. Brown, M. W. Covert, S. C. Doney, S. T. Dyhrman, A. S. Edison, A. M. Eren, N. M. Levine, L. Li, A. C. Ross, M. A. Saito, A. E. Santoro, D. Segrè, A. Shade, M. B. Sullivan, A. Vardi, Microbial metabolites in the marine carbon cycle. Nat. Microbiol. 7, 508– 523 (2022).

20. T. Tyrrell, A. Merico, “*Emiliania huxleyi*: bloom observations and the conditions that induce them” in *Coccolithophores* (Springer Berlin Heidelberg, Berlin, Heidelberg, 2004), pp. 75– 97.

21. P. M. Holligan, E. Fernández, J. Aiken, W. M. Balch, P. Boyd, P. H. Burkill, M. Finch, S. B. Groom, G. Malin, K. Muller, D. A. Purdie, C. Robinson, C. C. Trees, S. M. Turner, P. Van Der Wal, A biogeochemical study of the coccolithophore, *Emiliania huxleyi*, in the North Atlantic. Global Biogeochemical Cycles 7, 879–900 (1993).

22. U. Alcolombri, S. Ben-Dor, E. Feldmesser, Y. Levin, D. S. Tawfik, A. Vardi, Identification of the algal dimethyl sulfide-releasing enzyme: a missing link in the marine sulfur cycle. Science 348, 1466–1469 (2015).

23. W. H. Wilson, G. A. Tarran, D. Schroeder, M. Cox, J. Oke, G. Malin, Isolation of viruses responsible for the demise of an *Emiliania huxleyi* bloom in the English Channel. J. Mar. Biol. Assoc. U. K. 82, 369–377 (2002).

24. F. Vincent, U. Sheyn, Z. Porat, D. Schatz, A. Vardi, Visualizing active viral infection reveals diverse cell fates in synchronized algal bloom demise. Proc. Natl. Acad. Sci. U. S. A. 118, e2021586118 (2021).

25. G. Hevroni, F. Vincent, C. Ku, U. Sheyn, A. Vardi, Daily turnover of active giant virus infection during algal blooms revealed by single-cell transcriptomics. Sci. Adv. 9, eadf7971 (2023).

26. Y. Lehahn, I. Koren, D. Schatz, M. Frada, U. Sheyn, E. Boss, S. Efrati, Y. Rudich, M. Trainic, S. Sharoni, C. Laber, G. R. DiTullio, M. J. L. Coolen, A. M. Martins, B. A. S. Van Mooy, K. D. Bidle, A. Vardi, Decoupling physical from biological processes to assess the impact of viruses on a mesoscale algal bloom. Curr. Biol. 24, 2041–2046 (2014).

27. D. M. Morens, J. K. Taubenberger, A. S. Fauci, Predominant role of bacterial pneumonia as a cause of death in pandemic influenza: implications for pandemic influenza preparedness. J. Infect. Dis. 198, 962–970 (2008).

28. R. Sharan, A. N. Bucşan, S. Ganatra, M. Paiardini, M. Mohan, S. Mehra, S. A. Khader, D. Kaushal, Chronic immune activation in TB/HIV co-infection. Trends Microbiol. 28, 619–632 (2020).

29. L. Behrendt, U. Alcolombri, J. E. Hunter, S. Smriga, T. Mincer, D. P. Lowenstein, Y. Yawata, F. J. Peaudecerf, V. I. Fernandez, H. F. Fredricks, H. Almblad, J. J. Harrison, R. Stocker, B. A. S. Van Mooy, Microbial dietary preference and interactions affect the export of lipids to the deep ocean. Science 385, eaab2661 (2024).

30. S. Malitsky, C. Ziv, S. Rosenwasser, S. Zheng, D. Schatz, Z. Porat, S. Ben-Dor, A. Aharoni, A. Vardi, Viral infection of the marine alga *Emiliania huxleyi* triggers lipidome remodeling and induces the production of highly saturated triacylglycerol. New Phytol. 210, 88–96 (2016).

31. D. Schatz, S. Rosenwasser, S. Malitsky, S. G. Wolf, E. Feldmesser, A. Vardi, Communication via extracellular vesicles enhances viral infection of a cosmopolitan alga. Nat. Microbiol. 2, 1485–1492 (2017).

32. A. Vardi, L. Haramaty, B. A. S. Van Mooy, H. F. Fredricks, S. A. Kimmance, A. Larsen, K. D. Bidle, Host-virus dynamics and subcellular controls of cell fate in a natural coccolithophore population. Proc. Natl. Acad. Sci. U. S. A. 109, 19327–19332 (2012).

33. C. P. Laber, J. E. Hunter, F. Carvalho, J. R. Collins, E. J. Hunter, B. M. Schieler, E. Boss, K. More, M. Frada, K. Thamatrakoln, C. M. Brown, L. Haramaty, J. Ossolinski, H. Fredricks, J. I. Nissimov, R. Vandzura, U. Sheyn, Y. Lehahn, R. J. Chant, A. M. Martins, M. J. L. Coolen, A. Vardi, G. R. DiTullio, B. A. S. Van Mooy, K. D. Bidle, Coccolithovirus facilitation of carbon export in the North Atlantic. Nat. Microbiol. 3, 537–547 (2018).

34. B. S. Lambert, J.-B. Raina, V. I. Fernandez, C. Rinke, N. Siboni, F. Rubino, P. Hugenholtz, G. W. Tyson, J. R. Seymour, R. Stocker, A microfluidics-based in situ chemotaxis assay to study the behaviour of aquatic microbial communities. Nat. Microbiol. 2, 1344–1349 (2017).

35. E. Ivars-Martinez, A.-B. Martin-Cuadrado, G. D’Auria, A. Mira, S. Ferriera, J. Johnson, R. Friedman, F. Rodriguez-Valera, Comparative genomics of two ecotypes of the marine planktonic copiotroph *Alteromonas macleodii* suggests alternative lifestyles associated with different kinds of particulate organic matter. ISME J. 2, 1194–1212 (2008).

36. G. Cai, X. Yu, H. Wang, T. Zheng, F. Azam, Nutrient-dependent interactions between a marine copiotroph *Alteromonas* and a diatom *Thalassiosira pseudonana*. MBio 14, e0094023 (2023).

37. D. Wiener, Z. Bartolek, R. Dunklin, E. V. Armbrust, Growth phase-specific gene regulation and algicidal interactions between a new *A. macleodii* strain and the model diatom *T. pseudonana*. bioRxiv (2025).

38. R. J. Henshaw, J. Moon, M. R. Stehnach, B. P. Bowen, S. M. Kosina, T. R. Northen, J. S. Guasto, S. A. Floge, Metabolites from intact phage-infected *Synechococcus* chemotactically attract heterotrophic marine bacteria. Nat. Microbiol. 9, 3184–3195 (2024).

39. A. A. Shibl, A. Isaac, M. A. Ochsenkühn, A. Cárdenas, C. Fei, G. Behringer, M. Arnoux, N. Drou, M. P. Santos, K. C. Gunsalus, C. R. Voolstra, S. A. Amin, Diatom modulation of select bacteria through use of two unique secondary metabolites. Proc. Natl. Acad. Sci. U. S. A. 117, 27445–27455 (2020).

40. S. Rosenwasser, C. Ziv, S. G. van Creveld, A. Vardi, Virocell metabolism: metabolic innovations during host-virus interactions in the ocean. Trends Microbiol. 24, 821–832 (2016).

41. G. Schleyer, N. Shahaf, C. Ziv, Y. Dong, R. A. Meoded, E. J. N. Helfrich, D. Schatz, S. Rosenwasser, I. Rogachev, A. Aharoni, J. Piel, A. Vardi, In plaque-mass spectrometry imaging of a bloom-forming alga during viral infection reveals a metabolic shift towards odd-chain fatty acid lipids. Nat. Microbiol. 4, 527–538 (2019).

42. J. I. Nissimov, A. Pagarete, F. Ma, S. Cody, D. D. Dunigan, S. A. Kimmance, M. J. Allen, Coccolithoviruses: a review of cross-kingdom genomic thievery and metabolic thuggery. Viruses 9 (2017).

43. J. M. Robertson, E. A. Garza, A. K. M. Stubbusch, C. L. Dupont, T. Hwa, N. A. Held, Marine bacteria *Alteromonas* spp. require UDP-glucose-4-epimerase for aggregation and production of sticky exopolymer. MBio 15, e0003824 (2024).

44. T. J. Mayers, A. R. Bramucci, K. M. Yakimovich, R. J. Case, A bacterial pathogen displaying temperature-enhanced virulence of the microalga *Emiliania huxleyi*. Front. Microbiol. 7, 892 (2016).

45. N. Barak-Gavish, B. Dassa, C. Kuhlisch, I. Nussbaum, A. Brandis, G. Rosenberg, R. Avraham, A. Vardi, Bacterial lifestyle switch in response to algal metabolites. Elife 12, e84400 (2023).

46. E. Segev, T. P. Wyche, K. H. Kim, J. Petersen, C. Ellebrandt, H. Vlamakis, N. Barteneva, J. N. Paulson, L. Chai, J. Clardy, R. Kolter, Dynamic metabolic exchange governs a marine algal-bacterial interaction. Elife 5, e17473 (2016).

47. A. Fromm, D. Schatz, S. Ben-Dor, E. Feldmesser, A. Vardi, Complete genome sequence of *Emiliania huxleyi* virus strain M1, isolated from an induced *E. huxleyi* bloom in Bergen, Norway. Microbiol. Resour. Announc. 11, e0007122 (2022).

48. M. D. Keller, R. C. Selvin, W. Claus, R. R. L. Guillard, Media for the culture of oceanic ultraphytoplankton. J. Phycol. 23, 633–638 (1987).

49. A. E. Parada, D. M. Needham, J. A. Fuhrman, Every base matters: assessing small subunit rRNA primers for marine microbiomes with mock communities, time series and global field samples. Environ. Microbiol. 18, 1403–1414 (2016).

50. A. Apprill, S. McNally, R. Parsons, L. Weber, Minor revision to V4 region SSU rRNA 806R gene primer greatly increases detection of SAR11 bacterioplankton. Aquat. Microb. Ecol. 75, 129–137 (2015).

51. B. J. Callahan, P. J. McMurdie, M. J. Rosen, A. W. Han, A. J. A. Johnson, S. P. Holmes, DADA2: High-resolution sample inference from Illumina amplicon data. Nat. Methods 13, 581–583 (2016).

52. C. Quast, E. Pruesse, P. Yilmaz, J. Gerken, T. Schweer, P. Yarza, J. Peplies, F. O. Glöckner, The SILVA ribosomal RNA gene database project: improved data processing and web-based tools. Nucleic Acids Res. 41, D590–6 (2013).

53. P. J. McMurdie, S. Holmes, phyloseq: an R package for reproducible interactive analysis and graphics of microbiome census data. PLoS One 8, e61217 (2013).

54. D. J. Lane, “16S/23S rRNA sequencing” in Nucleic Acid Techniques in Bacterial Systematics, E. Stackebrandt, M. Goodfellow, Eds. (John Wiley and Sons, New York, NY, 1991), pp. 115–175.

55. W. G. Weisburg, S. M. Barns, D. A. Pelletier, D. J. Lane, 16S ribosomal DNA amplification for phylogenetic study. J. Bacteriol. 173, 697–703 (1991).

56. E. Pruesse, J. Peplies, F. O. Glöckner, SINA: Accurate high-throughput multiple sequence alignment of ribosomal RNA genes. Bioinformatics 28, 1823–1829 (2012).

57. C. Ku, N. Barak-Gavish, M. Maienschein-Cline, S. J. Green, A. Vardi, Complete genome sequence of *Sulfitobacter* sp. strain D7, a virulent bacterium isolated from an *Emiliania huxleyi* algal bloom in the North Atlantic. Microbiol. Resour. Announc. 7 (2018).

58. F. Salvà Serra, F. Salvà-Serra, L. Svensson-Stadler, A. Busquets, D. Jaén-Luchoro, R. Karlsson, E. R. B. Moore, M. Gomila, A protocol for extraction and purification of high-quality and quantity bacterial DNA applicable for genome sequencing: a modified version of the Marmur procedure. Protoc. Exch., doi: 10.1038/protex.2018.084 (2018).

59. J. Marmur, A procedure for the isolation of deoxyribonucleic acid from micro-organisms. J. Mol. Biol. 3, 208–IN1 (1961).

60. P.-A. Chaumeil, A. J. Mussig, P. Hugenholtz, D. H. Parks, GTDB-Tk: a toolkit to classify genomes with the Genome Taxonomy Database. Bioinformatics 36, 1925–1927 (2019).

61. J. G. Caporaso, C. L. Lauber, W. A. Walters, D. Berg-Lyons, C. A. Lozupone, P. J. Turnbaugh, N. Fierer, R. Knight, Global patterns of 16S rRNA diversity at a depth of millions of sequences per sample. Proc. Natl. Acad. Sci. U. S. A. 108 Suppl 1, 4516–4522 (2011).

62. E. E. Clerc, J.-B. Raina, J. M. Keegstra, Z. Landry, S. Pontrelli, U. Alcolombri, B. S. Lambert, V. Anelli, F. Vincent, M. Masdeu-Navarro, A. Sichert, F. De Schaetzen, U. Sauer, R. Simó, J.-H. Hehemann, A. Vardi, J. R. Seymour, R. Stocker, Strong chemotaxis by marine bacteria towards polysaccharides is enhanced by the abundant organosulfur compound DMSP. Nat. Commun. 14, 8080 (2023).

63. R. Stocker, J. R. Seymour, Ecology and physics of bacterial chemotaxis in the ocean. Microbiol. Mol. Biol. Rev. 76, 792–812 (2012).

64. E. E. Clerc, J.-B. Raina, B. S. Lambert, J. Seymour, R. Stocker, In situ chemotaxis assay to examine microbial behavior in aquatic ecosystems. J. Vis. Exp. 159, e61062 (2020).

65. A. R. Bramucci, A. Focardi, C. Rinke, P. Hugenholtz, G. W. Tyson, J. R. Seymour, J.-B. Raina, Microvolume DNA extraction methods for microscale amplicon and metagenomic studies. ISME Commun. 1, 79 (2021).

66. C. Rinke, S. Low, B. J. Woodcroft, J.-B. Raina, A. Skarshewski, X. H. Le, M. K. Butler, R. Stocker, J. Seymour, G. W. Tyson, P. Hugenholtz, Validation of picogram- and femtogram-input DNA libraries for microscale metagenomics. PeerJ 4, e2486 (2016).

67. T. Magoč, S. L. Salzberg, FLASH: fast length adjustment of short reads to improve genome assemblies. Bioinformatics 27, 2957–2963 (2011).

68. M. Martin, Cutadapt removes adapter sequences from high-throughput sequencing reads. EMBnet J. 17, 10 (2011).

69. F. Vincent, G. Schleyer, C. Kuhlisch, C. Marrasé, R. Simó, J. Egge, A. Vardi, D. Schatz, AQUACOSM VIMS-Ehux – Core data. Dryad (2020).

70. E. P. Ivanova, S. Flavier, R. Christen, Phylogenetic relationships among marine *Alteromonas*-like proteobacteria: emended description of the family *Alteromonadaceae* and proposal of *Pseudoalteromonadaceae* fam. nov., *Colwelliaceae* fam. nov., *Shewanellaceae* fam. nov., *Moritellaceae* fam. nov., *Ferrimonadaceae* fam. nov., Idiomarinaceae fam. nov. and Psychromonadaceae fam. nov. Int. J. Syst. Evol. Microbiol. 54, 1773–1788 (2004).

