## Supplementary Information for "Synergy in viral-bacterial coinfection expedites algal bloom demise"

Figures S1 to S13

Legends for Tables S1 to S6

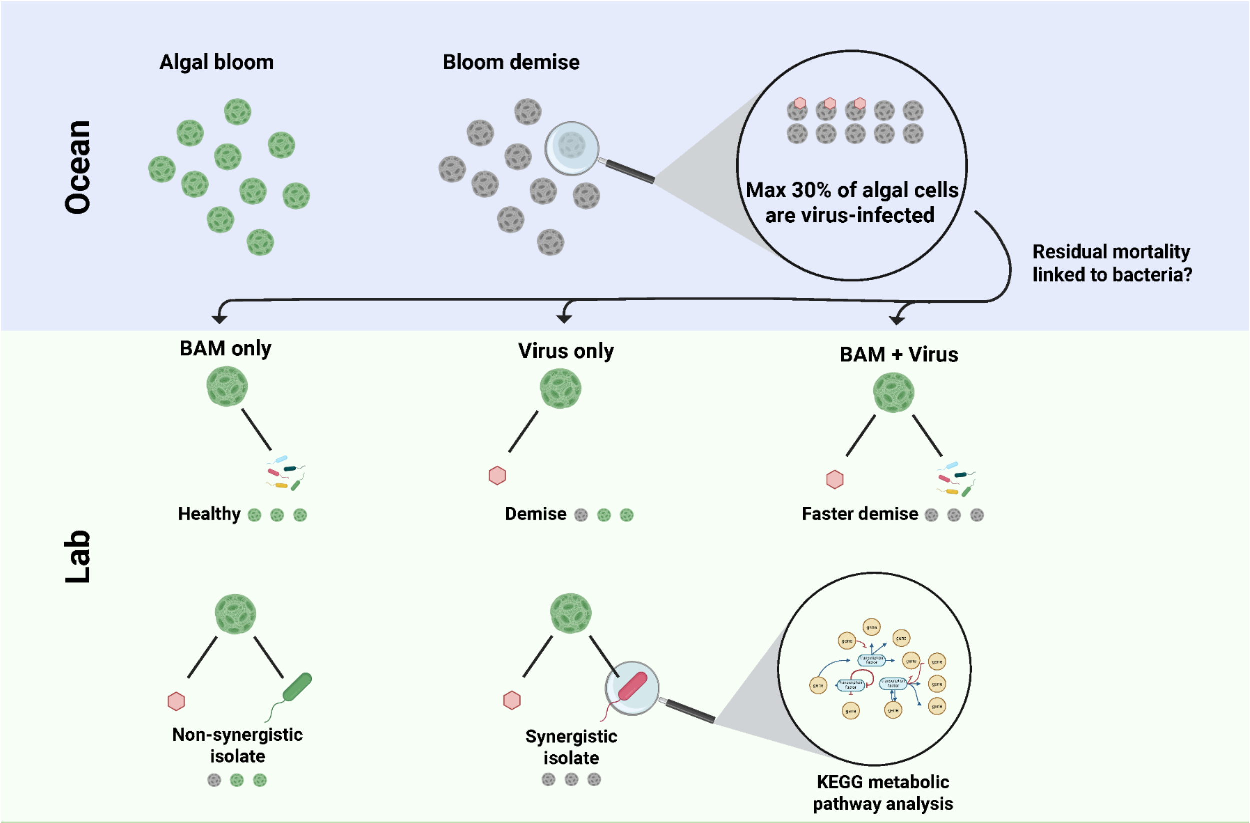


**Figure S1. Developing a tripartite alga-virus-bacteria model system to recapitulate microbial interactions during algal bloom demise.** Single cell analysis of viral gene expression during *G. huxleyi* bloom demise revealed that at maximum 30% of the population were virus-infected (*18*). The residual cell mortality supporting a synchronized algal bloom demise might be linked to induced pathogenicity in the algal bloom microbiome. To study algae-virus-bacteria interactions, we isolated from the bloom and demise phases *G. huxleyi*, EhV and the bloom-associated microbiome (BAM) and followed the dynamics of algal cultures following individual and co-infections with BAM and the virus. Synergistic bacteria that enhanced population demise were isolated and their metabolic niche evaluated via functional pathway analysis.

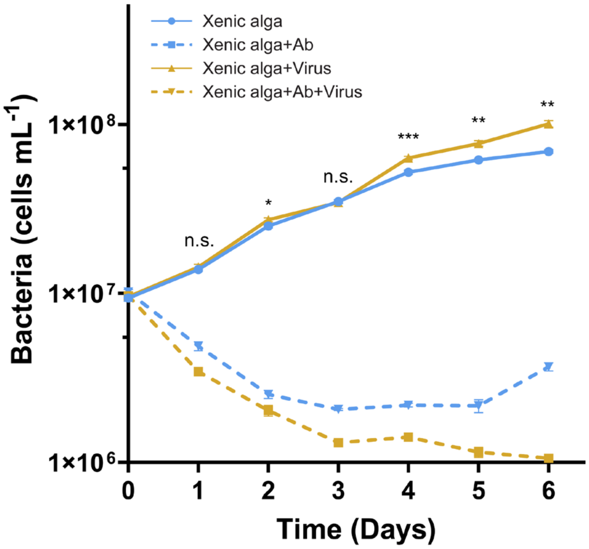


**Figure S2. Antibiotics decrease bacterial abundances in xenic algal cultures.** Bacterial abundance in cultures of *G. huxleyi* under four conditions: uninfected (blue lines) and EhV-infected (yellow lines) with (dashed lines) and without (solid lines) antibiotics (Ab). n.s. - not significant, * $p < 0.05$, ** $p < 0.01$, *** $p < 0.001$, **** $p < 10^{-4}$; statistical significance determined by two-way ANOVA with Sídák's post-hoc test; n = 4.

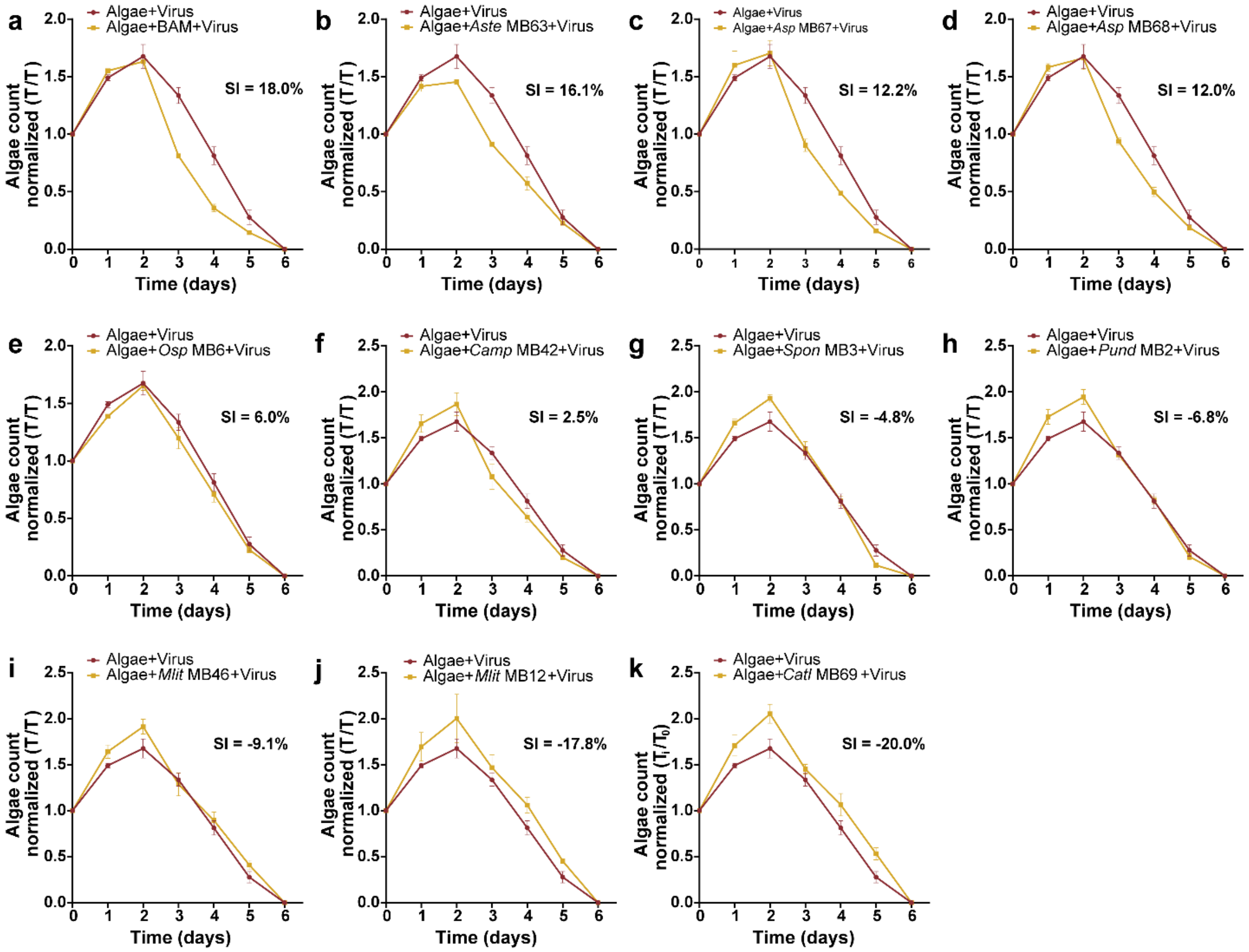


**Figure S3. Synergy indices (SIs) of BAM-derived bacteria.** Abundances of *G. huxleyi* following viral infection in the presence (yellow) or absence (red) of (a) the bloom-associated microbiome (BAM) and of (b-k) ten bacteria isolated from the BAM (see **Table S1** for strain information). SI values calculated from the abundance data are inserted in bold text. n = 3.

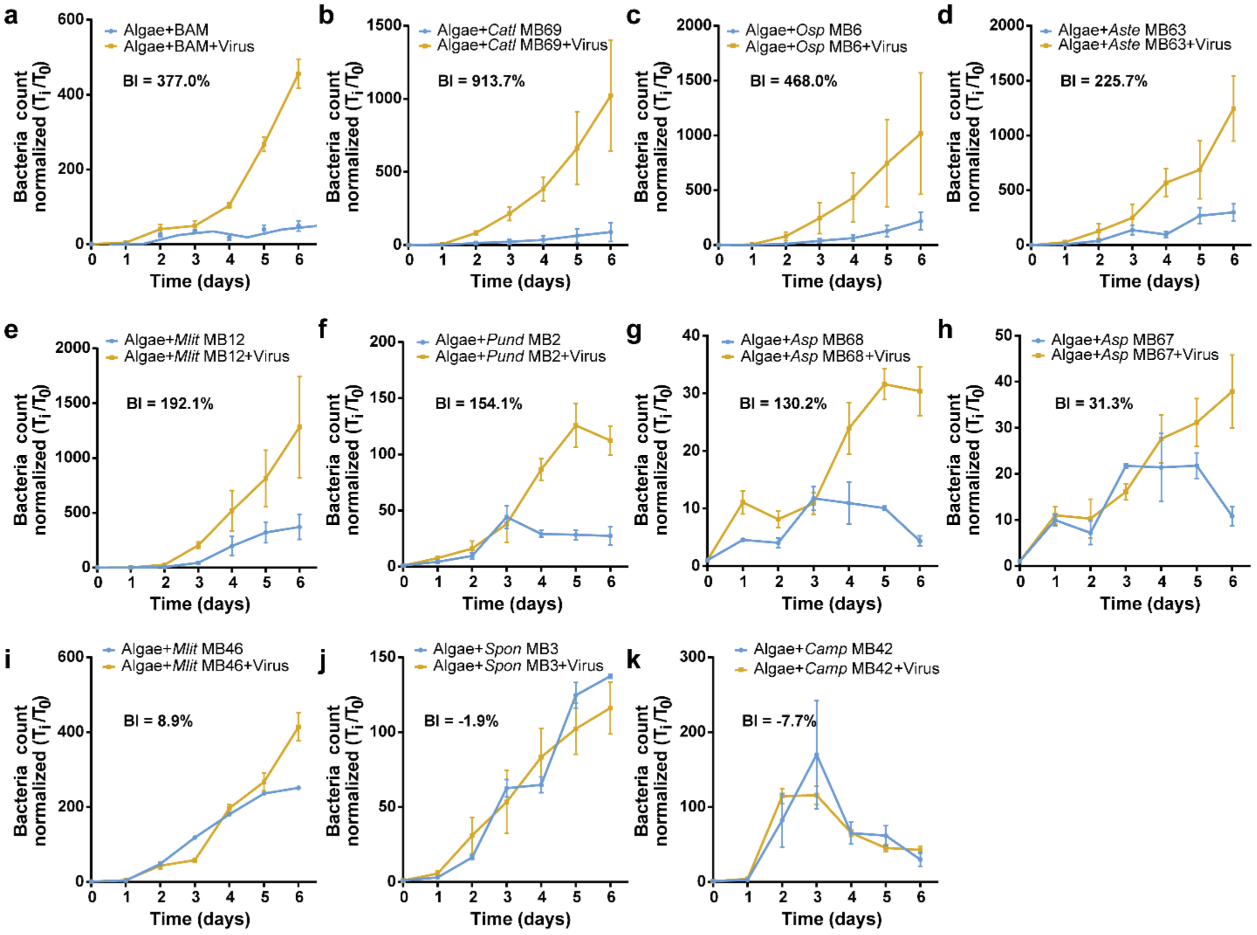


**Figure S4. Benefit indices (BIs) of BAM-derived bacteria.** Abundances of bacterial isolates from the bloom-associated microbiome (BAM) in co-culture with *G. huxleyi* and in the presence (yellow) or absence (blue) of viral infection. The growth dynamics of (a) the BAM and of (b-k) ten BAM-derived bacterial strains (see **Table S1** for strain information). BI values calculated from the abundance data are inserted in bold text. n = 3.

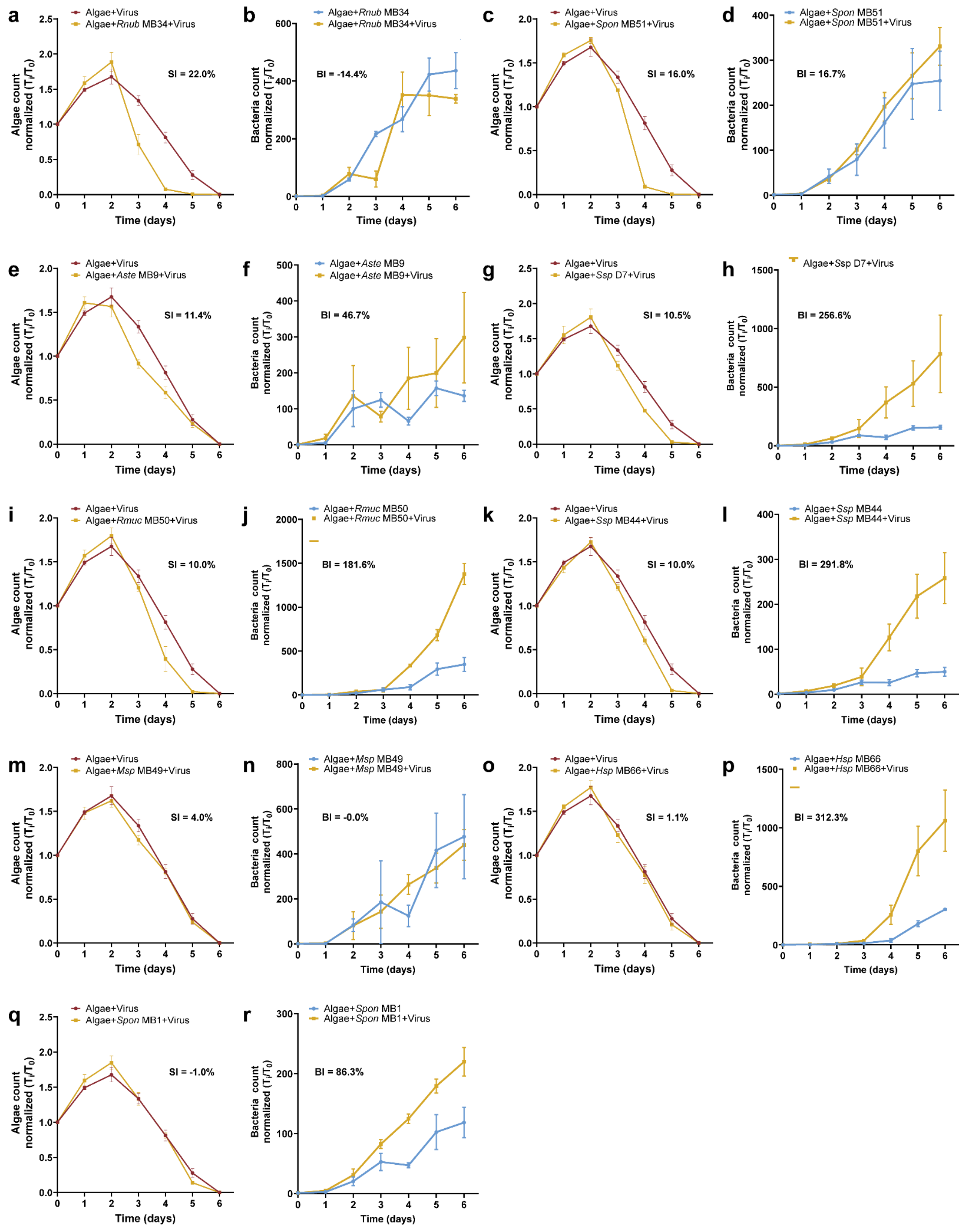


**Figure S5. Synergy indices (SIs) and benefit indices (BIs) of mesocosm-derived bacteria.** Abundances of *G. huxleyi* following EhV infection in the presence (yellow) or absence (red) of nine bacteria isolated from an induced mesocosm *G. huxleyi* bloom. Bacterial abundances of nine bacteria isolated from an induced mesocosm bloom in co-culture with *G. huxleyi* and in the presence (yellow) or absence (blue) of viral infection. SI and BI values calculated from the abundance data are inserted in bold text. See **Table S1** for strain information. n = 3.

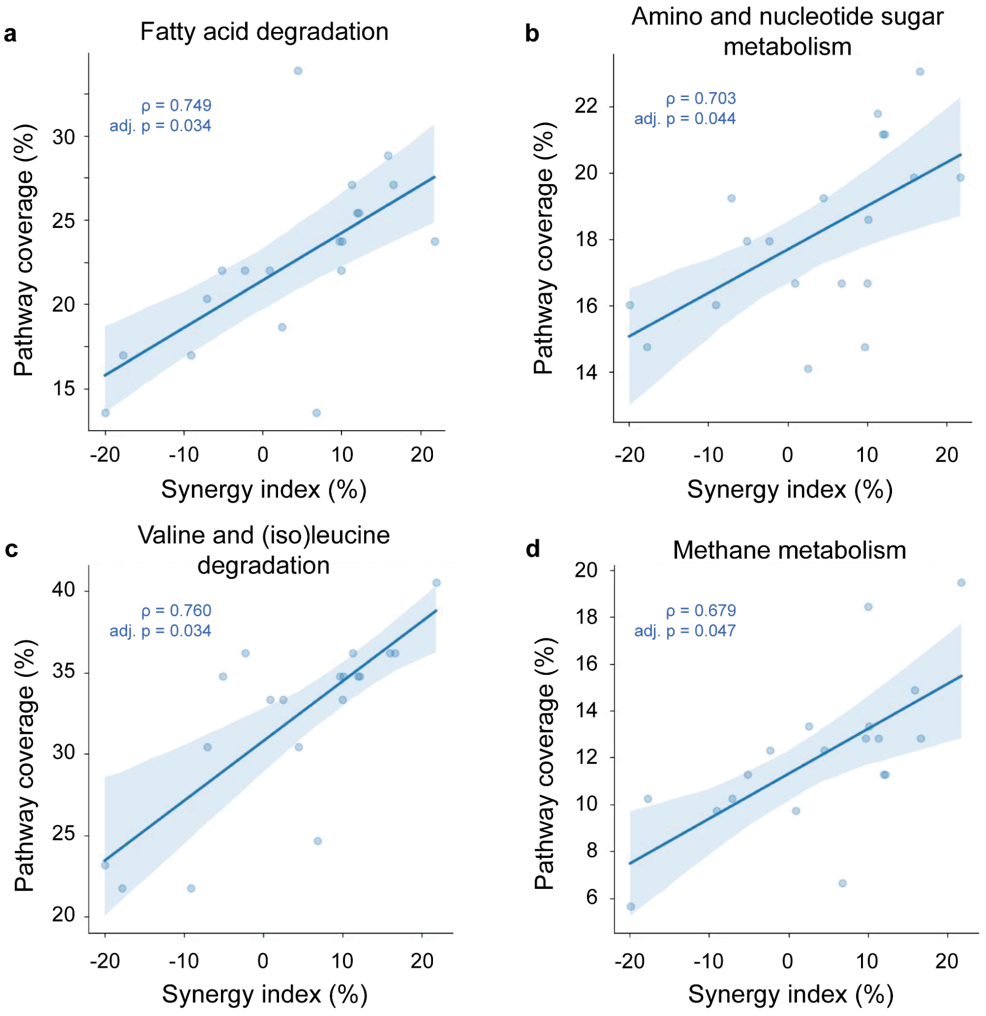


**Figure S6. Correlation of the bacterial SI with the completeness of KEGG metabolic pathways.** Significantly positive correlations were found between the SI and the completeness of the (a) fatty acid degradation pathway (#00071), (b) amino sugar and nucleotide sugar metabolism pathway (#00520), (c) valine, leucine and (iso)leucine degradation pathway (#00280), and (d) methane metabolism pathway (#00680) across nineteen *G. huxleyi* bloom-derived bacteria. Shaded areas represent the 95% confidence interval of the linear regression. See **Tables S2-S5** for gene information for each pathway.

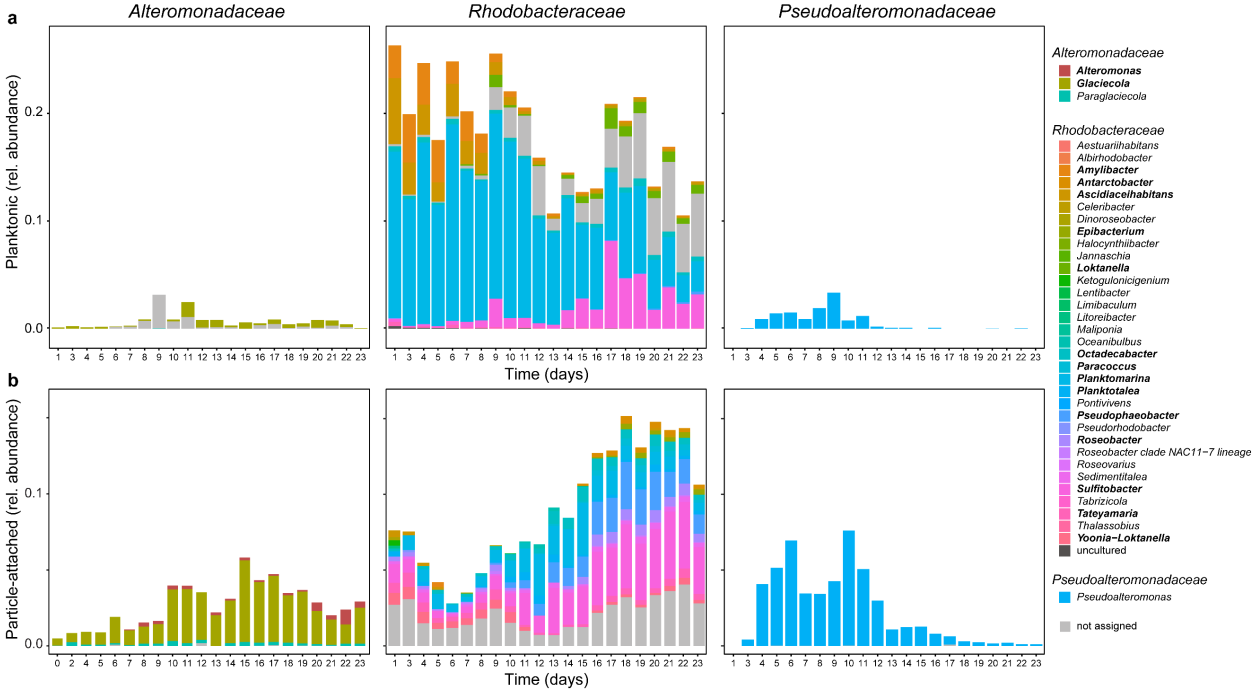


**Figure S7. Taxonomic composition of planktonic and particle-attached bacteria during *G. huxleyi* bloom succession using 16S rRNA amplicon sequencing.** Genus level composition is shown for (a) the planktonic size fraction (<2 µm and >0.2 µm; data modified from (*18*) and for (b) the particle-attached size fraction (<200 µm and >20 µm) of selected bacterial families. The *Alteromonadaceae* family was dominated by members of the genus *Glaciecola*, which is highly similar to the genus *Alteromonas* on the level of the 16S rRNA gene sequence (*70*). Planktonic *Rhodobacteraceae* decreased over time and shifted from a high relative abundance in e.g. *Planktomarina* and *Amylibacter* to *Sulfitobacter* and *Loktanella*, while the particle-attached fraction increased in relative abundance with a dominance in e.g. *Planktomarina, Sulfitobacter* and *Pseudophaeobacter*. Both planktonic and particle-attached *Pseudoalteromonadaceae* decreased over time and were constituted only of *Pseudalteromonas* members. Dominant genera are indicated in bold letters in the legend.

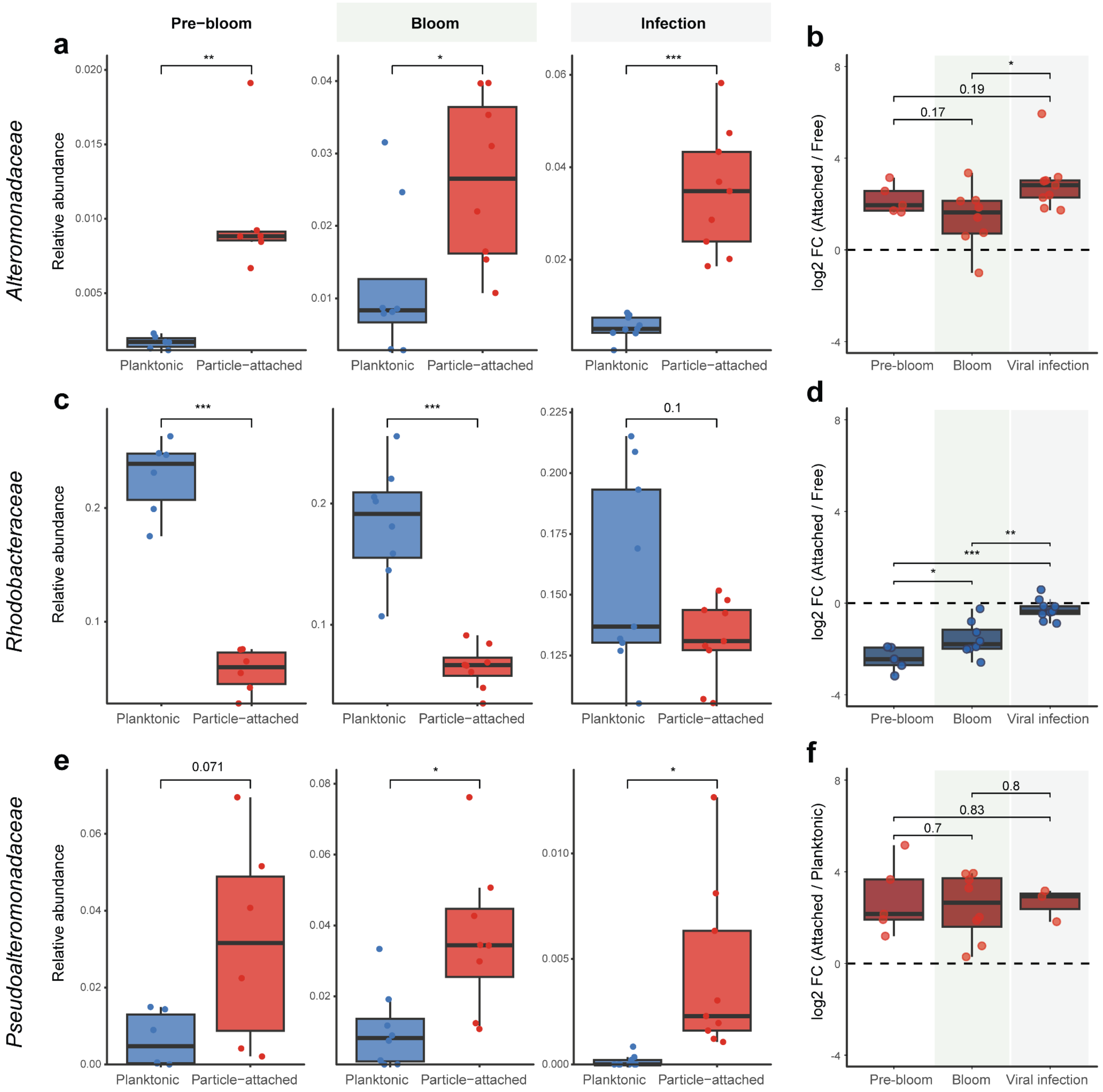


**Figure S8. Occurrences of *Alteromonadaceae*, *Rhodobacteraceae* and *Pseudoalteromonadaceae* in planktonic and particle-attached size fractions during *G. huxleyi* bloom succession.** Relative abundances of *Alteromonadaceae* (a), *Rhodobacteraceae* (c) and *Pseudoalteromonadaceae* (e) in the planktonic size fraction (<2 µm and >0.2 µm; data modified from (*18*)) and particle-attached size fraction (<200 µm and >20 µm) during the pre-bloom phase (days 0-6), bloom phase of *G. huxleyi* (days 7-14) and upon viral infection (days 15-23) using 16S rRNA amplicon sequencing. Log2 fold change (FC) of particle-attached to planktonic occurrences visualizes the enrichment of *Alteromonadaceae* (b) and *Pseudoalteromonadaceae* (f) in the particle-attached size fraction throughout the bloom succession, while *Rhodobacteraceae* (d) were more abundant in the planktonic size fraction. * $p < 0.05$, ** $p < 0.01$, *** $p < 0.001$; statistical significance of relative abundances was determined by Kruskal-Wallis tests and of log2 FC by pairwise Welch t-tests.

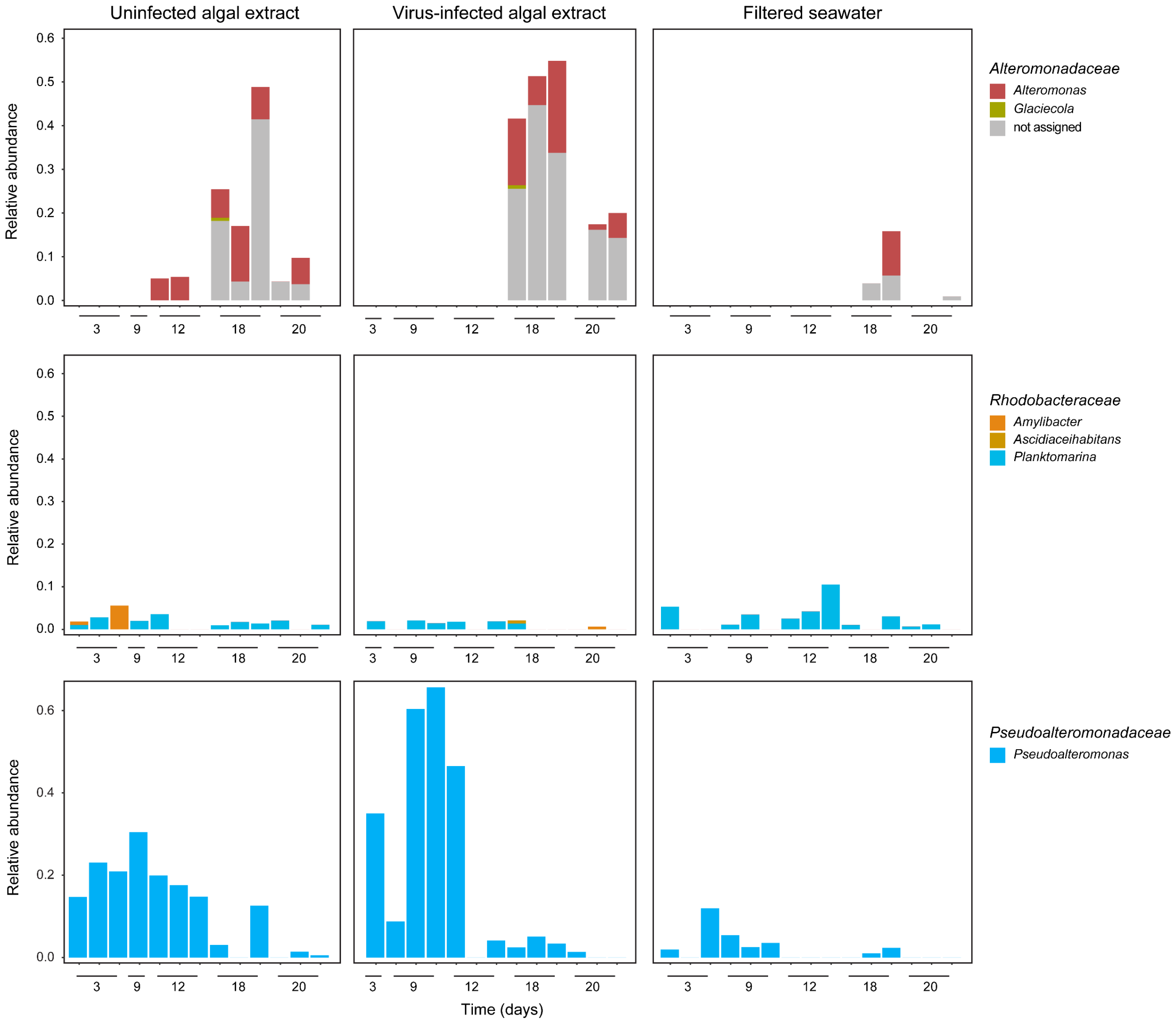


**Figure S9. Taxonomic composition of chemotactic bacteria during *G. huxleyi* bloom succession using 16S rRNA amplicon sequencing.** *In-situ* chemotaxis assays (ISCA) were performed at selected days throughout phytoplankton bloom succession with the bloom associated bacteria using two different *G. huxleyi* extracts, namely cell extracts derived from uninfected and virus-infected *G. huxleyi* CCMP 2090 cultures. Filtered seawater was used as a reference. Genus level composition is shown for *Alteromonadaceae* that exhibited the strongest chemoattraction to extracts from infected cells during the viral infection phase of the *G. huxleyi* bloom (days 15-23), for *Rhodobacteraceae* that showed minimal chemoattraction throughout the entire phytoplankton bloom succession, and for *Pseudoalteromonadaceae* that demonstrated the strongest chemoattraction during the pre-bloom phase (days 0-14).

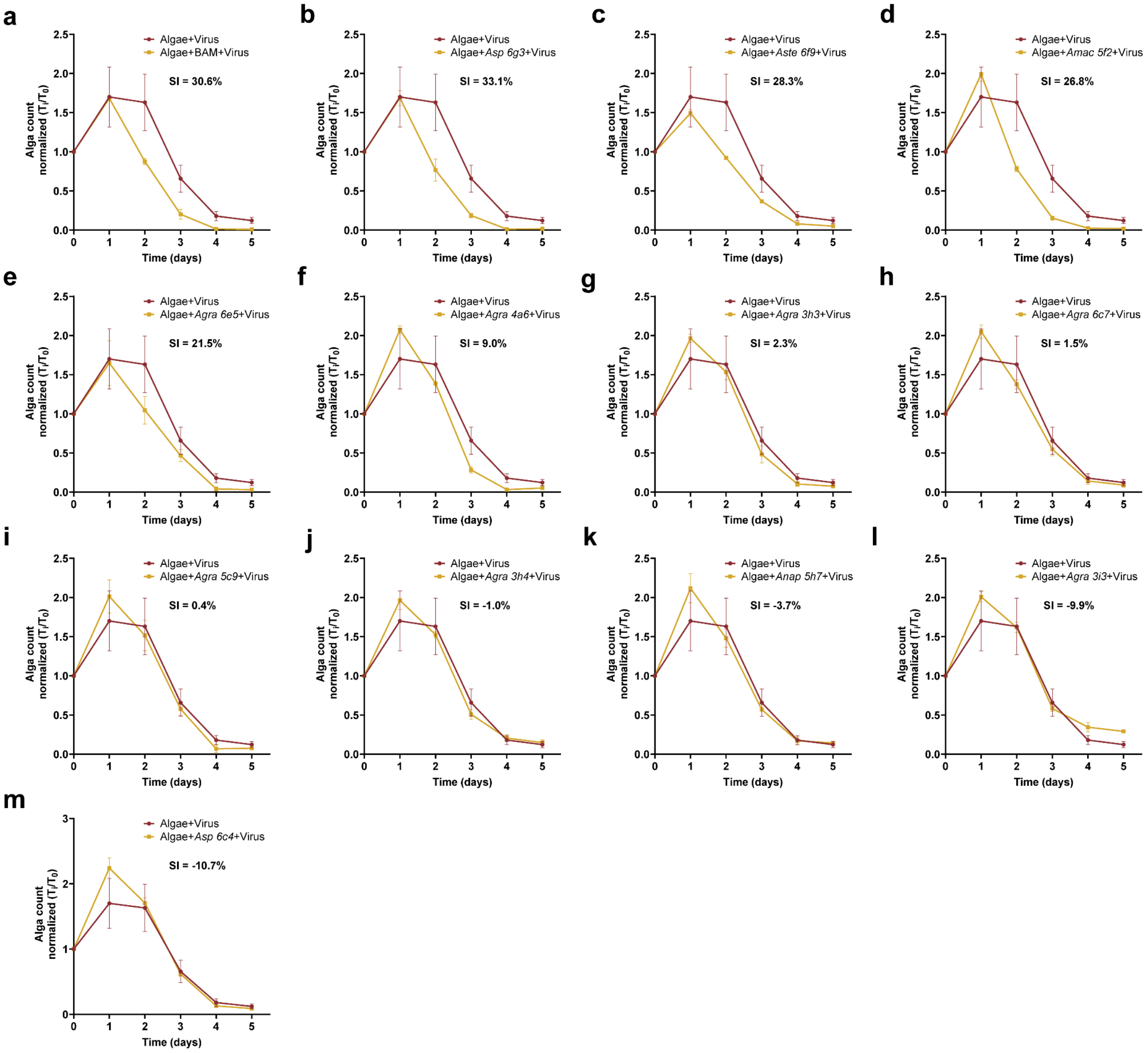


**Figure S10. Synergy indices (SIs) of chemotactic *Alteromonas* isolates.** Abundances of *G. huxleyi* following EhV infection in the presence (yellow) or absence (red) of chemotactic *Alteromonas* strains. The growth dynamics of (a) the bloom-associated microbiome (BAM) and of (b-m) twelve chemotactic *Alteromonas* strains isolated from the ISCA experiment are shown (see **Table S1** for strain information). n = 3.

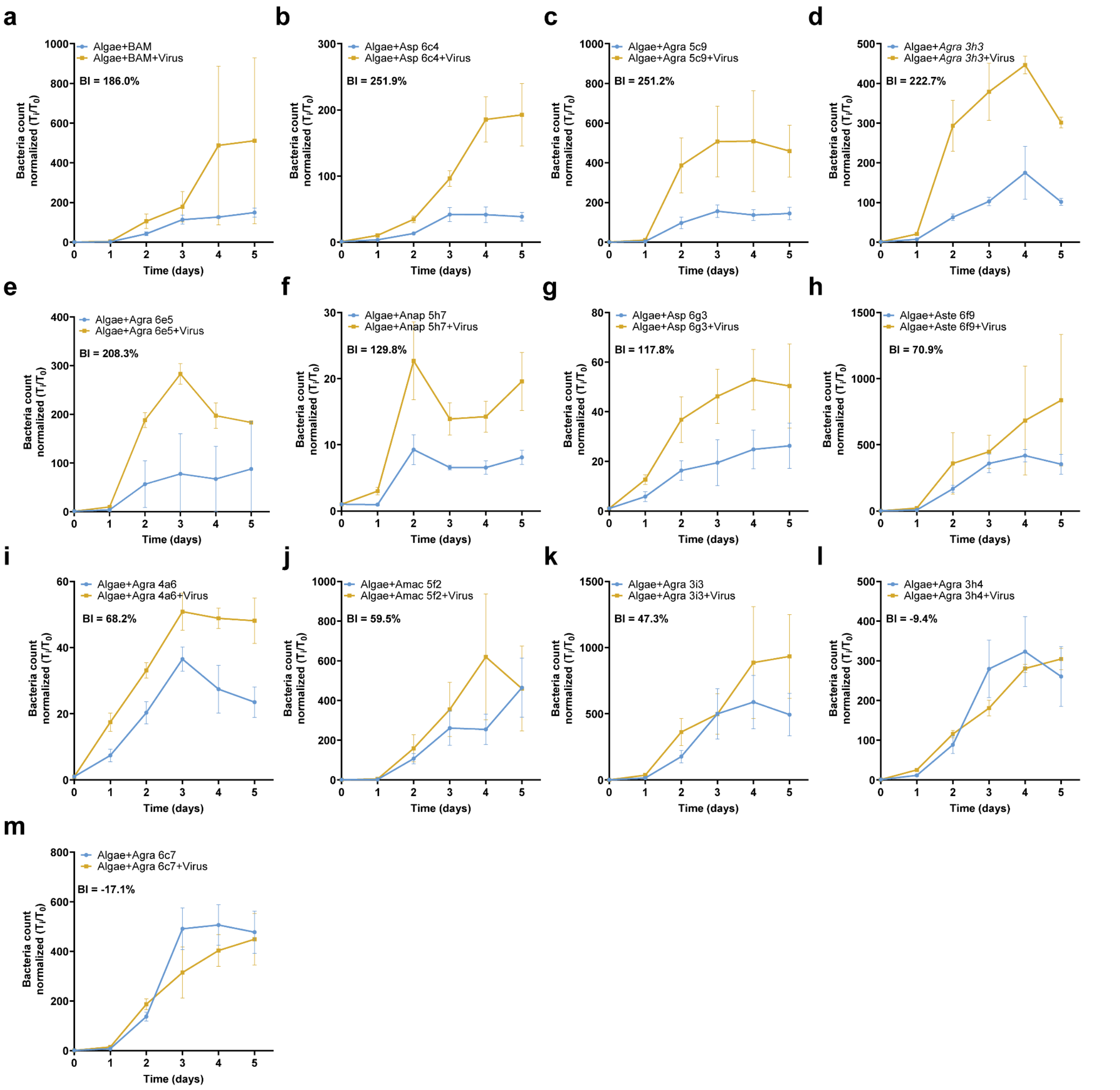


**Figure S11. Benefit indices (BIs) of chemotactic *Alteromonas* isolates.** Bacterial abundances within *G. huxleyi* co-cultures in the presence (yellow) or absence (blue) of viral infection. The growth dynamics of (a) the bloom-associated microbiome (BAM) and of (b-m) twelve chemotactic *Alteromonas* strains isolated from the ISCA experiment (see **Table S1** for strain information) are shown. n = 3.

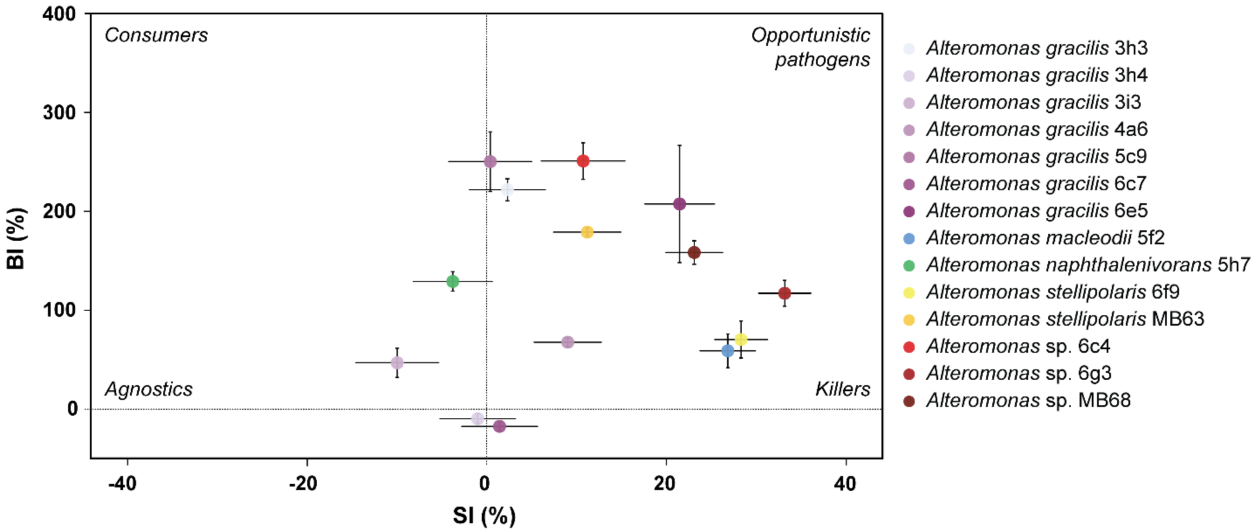


**Figure S12. Ecological strategies for algal bloom-associated *Alteromonas* based on SI and BI.** Synergy indices (SI) and benefit indices (BI) of two *Alteromonas* strains that were isolated from BAM7 (*Alteromonas* sp. MB63 and *Alteromonas* sp. MB68) and twelve *Alteromonas* strains that were isolated upon chemotaxis towards algal cell extracts. n = 3.

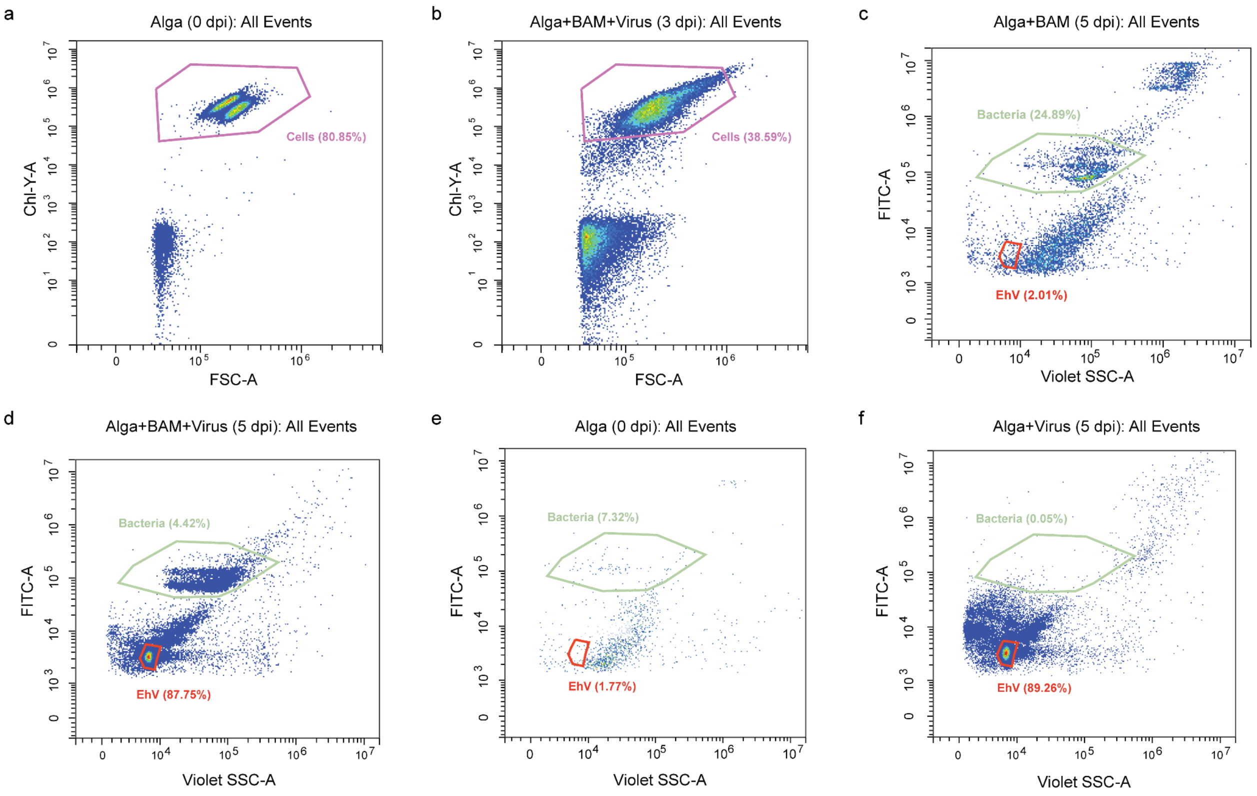


**Figure S13. Flow cytometry images visualizing the gating strategy for tripartite interaction experiments.** Uninfected (a, c and e) and virus-infected (b, d and f) cultures of axenic *G. huxleyi* RCC6946 with (b-d) and without (a, e and f) inoculation of a bloom-associated microbiome (BAM) were analyzed by flow cytometry. Algal cells were enumerated by plotting the autofluorescence of chlorophyll (em: 663-737 nm) versus forward scatter (a and b; gate "Cells"). Bacteria (gate "Bacteria") and viruses (c-f; gate "EhV) were enumerated after DNA staining by plotting fluorescence (ex.: 488 nm, em.: 500-550 nm) versus side scatter (ex.: 405 nm). The gating of algal cells was based on the cell population of uninfected cultures at day 0 of each experiment. The bimodal cell distribution visible in (a) corresponds to calcified and non-calcified subpopulations. The gating of bacteria and viruses was based on the population of virus-infected and BAM-inoculated cultures at the end of each experiment. The highest fluorescent particles in (c) correspond to stained algal cells.

### Supplementary Table legends

**Table S1. Summary of bacterial strains derived from an induced *G. huxleyi* mesocosm bloom in 2018.** Metadata for 31 bacteria tested for their synergy during virus-induced algal demise, including the origin of isolation, genome sequencing method, genome size (in bp), number of contigs, GC content (in %), genome completeness and contamination (in %), taxonomic annotation using GTDB-Tk with the Average nucleotide identity (ANI) and alignment fraction (AF) scores to the closest reference genome, the NCBI BioProject and accession numbers, as well as the reference of their first published description. *Sulfitobacter* sp. D7 was included as a reference as reported algal pathogen derived from a coccolithophore bloom in the North Atlantic.

**Table S2-S5. Occurrence of SI-correlated metabolic pathway genes across bacterial strains.** Summary of gene homologies for four SI-correlated KEGG pathways, namely the fatty acid degradation (map #00071; **Table S2**), amino sugar and nucleotide sugar metabolism (map #00520; **Table S3**), branched-chain amino acid degradation (map #00280; **Table S4**), and methane metabolism pathway (map #00680; **Table S5**) across all isolated bacterial strains (n = 19), reported as –log10-transformed e-values (higher values indicate stronger homology). Genes were annotated using PFAM. n.a. = not available.

**Table S6. Composition of modified K/2 medium.** The medium composition of this modified K/2 medium follows the modifications by Ian Probert provided by the Roscoff Culture Collection (RCC) with the addition of $CuSO_4 \cdot 5\,H_2O$ as in the original recipe for K/2 (*48*).
